# Massively parallel assessment of human variants with base editor screens

**DOI:** 10.1101/2020.05.17.100818

**Authors:** Ruth E Hanna, Mudra Hegde, Christian R Fagre, Peter C DeWeirdt, Annabel K Sangree, Zsofia Szegletes, Audrey Griffith, Marissa N Feeley, Kendall R Sanson, Yossef Baidi, Luke W Koblan, David R Liu, James T Neal, John G Doench

## Abstract

Understanding the functional consequences of single-nucleotide variants is critical to uncovering the genetic underpinnings of diseases, but technologies to characterize variants are limiting. Here we leverage CRISPR-Cas9 cytosine base editors in pooled screens to scalably assay variants at endogenous loci in mammalian cells. We benchmark the performance of base editors in positive and negative selection screens and identify known loss-of-function mutations in *BRCA1* and *BRCA2* with high precision. To demonstrate the utility of base editor screens to probe small molecule-protein interactions, we conduct screens with BH3 mimetics and PARP inhibitors and identify point mutations that confer drug sensitivity or resistance. Finally, we create a library of 52,034 clinically-observed variants in 3,584 genes and conduct screens in the presence of cellular stressors, identifying loss-of-function variants in numerous DNA damage repair genes. We anticipate that this screening approach will be broadly useful to readily and scalably functionalize genetic variants.

## INTRODUCTION

A major challenge in genomics is the functional characterization of precise genetic variants at a large scale. Although genome-wide association studies (GWAS) have identified tens of thousands of associations between single nucleotide polymorphisms (SNPs) and phenotypes, identification of the causal variants lags behind. Ascertaining the functional consequence of a causal variant is more difficult still, typically requiring low-throughput genome editing to introduce the variant and characterize its functional significance. Functional characterization of genetic variants is also a bottleneck for rare disease research and cancer genomics; sequencing of clinical isolates in both contexts often uncovers variants that remain untested for their functional consequence, further expanding the list of variants of uncertain significance (VUS).

Many technologies for variant screening – sometimes called multiplexed assays of variant effects, or MAVEs (Starita et al., 2017; Weile et al., 2017) – offer different strengths and weaknesses. One general category of MAVEs are assays in which a predefined set of variants is introduced and functionally screened. For example, saturation mutagenesis (Melnikov et al., 2014; Patwardhan et al., 2009) can be used to screen all possible single-nucleotide variants in coding or non-coding regions, but relies on exogenous overexpression of the variant of interest, which may not always phenocopy endogenous variants, and is limited by the size of the cassette that can be introduced, making it difficult to assay large genes. Saturation genome editing (Findlay et al., 2014, 2018), which relies on homology-directed repair (HDR), has been used to screen all possible variants in several exons of *BRCA1*; while HDR approaches are effective in yeast (Sharon et al., 2018), the low efficiency of HDR in human cells has restricted its use to rare near-haploid lines.

The suite of CRISPR-based tools offers several options to mutagenize loci. Cas9 nuclease results in indels of varying lengths in most cell types, and several groups have used this approach to identify mutations that confer drug resistance to many families of small molecules (Donovan et al., 2017; Neggers et al., 2018; Pettitt et al., 2018; Vinyard et al., 2019). Although this strategy is readily deployable, wild-type Cas9 introduces a substantial fraction of out-of-frame indels that result in protein truncations and therefore non-functional protein variants, limiting its usefulness for mutagenesis. An alternative is to use catalytically-deactivated Cas9 (dCas9) to recruit a highly mutagenic agent to semi-randomly introduce substitutions at endogenous loci in mammalian cells (Hess et al., 2016; Ma et al., 2016). While these approaches do not directly generate out-of-frame indels, the low efficiency and heterogeneity of the substitutions limits their use in negative selection assays.

CRISPR base editors (Gaudelli et al., 2017; Komor et al., 2016), which introduce transition mutations at target loci, are an attractive candidate for sub-saturating variant screening for several reasons. First, they require only a single guide RNA (sgRNA) to direct the base editor to the desired locus; therefore, they can be used to screen large genes or genomic regions that could not easily be assayed by saturation mutagenesis. Second, unlike hyperactive editors, base editors edit in a relatively predictable fashion, generally editing all target nucleotides in a defined window with minimal indels (Komor et al., 2016). These properties should make base editors effective when used for negative selection screens, which require that cells receiving the same sgRNA generally receive the same edit. Moreover, base editors function in post-mitotic cells that do not generally support HDR (Levy et al., 2020; Rees and Liu, 2018; Yeh et al., 2018). The programmable and predictable nature of base editors also means that the results of a base editing screen can be initially assessed by sequencing the sgRNA only (rather than the full genomic region), followed by targeted sequencing of hit loci from the primary screen to identify individual causal variants. This reduces sequencing costs and allows for larger libraries that can be used to interrogate many disparate genomic loci at once.

Initial screens with base editor technology in mammalian cells have introduced stop codons (Kuscu et al., 2017) or tiled all possible sgRNAs across a small number of genes (Jun et al., 2020; Kweon et al., 2019), and such screens have recently been applied at scale in yeast (Despres et al., 2020). A comprehensive demonstration of the utility of base editor screens would facilitate widespread uptake of such screens as an approach to assay human genetic variants. Here, we present data from 25 pooled screens with cytosine base editors in human cell lines. We test two variants of base editors and demonstrate that the higher-performing variant effectively identifies loss-of-function mutations in positive and negative selection assays. We then use this base editor screening toolkit to interrogate variants in *BRCA1* and *BRCA2*, conduct modifier screens with 7 small molecules, and screen 52,034 clinically-observed genetic variants across several growth conditions. Our results highlight that base editor screens are widely useful to functionally characterize variants in a scalable, pooled fashion.

## RESULTS

### Benchmarking base editors in negative and positive selection screens

To assess the feasibility of performing pooled screens with base editors, we used a previously-described library (Sanson et al., 2019), which included all possible *S. pyogenes* sgRNAs targeting the exons of 47 genes that, when inactivated, confer a phenotype that is readily assayed in a pooled viability screen. These genes included 10 “pan-lethal” (essential) genes that are broadly required for cell survival, 4 genes whose knockout confers resistance to the BRAF-inhibitor vemurafenib (Shalem et al., 2014), as well as targeting and non-targeting controls, for a total library size of 12,141 sgRNAs. We cloned this library into lentiviral vectors that both express the sgRNA and contain either BE3.9max or BE4max (Koblan et al., 2018) (**Figure S1A**), which are C>T base editors that contain 1 and 2 copies of the uracil glycosylase inhibitor (UGI), respectively. As a control, we also cloned the library into a guide-only vector for use with cell lines stably expressing wild-type Cas9 (wtCas9).

We screened this library in A375 (melanoma) cells in both a negative selection assay (viability) and a positive selection assay (vemurafenib resistance). Unmodified parental cells (for base editor screens) or Cas9-expressing cells (for wtCas9 screens) were transduced with the library in duplicate, selected with puromycin for 6 days to eliminate uninfected cells, divided into 2 arms (dropout and vemurafenib), and cultured for 2 weeks at a representation of approximately 1,000 cells per sgRNA (**Figure 1A**). Genomic DNA was isolated at the final timepoint, sgRNAs were amplified by PCR, and sequenced. Log-fold changes of the dropout (no drug) arms were well correlated across replicates (Pearson r = 0.82 for BE3.9max; Pearson r = 0.79 for BE4max). We therefore merged replicates and calculated z-scores relative to targeting controls (guides tiling 4 cell surface marker genes; n = 671 for BE3.9max; n = 669 for BE4max). Next, we annotated sgRNAs with their predicted editing consequences using a simple heuristic: all C’s in positions 4-8 of the sgRNA were considered to be edited unless they were preceded by a G, as this motif is known to be highly disfavored by the APOBEC1 domain (Kluesner et al., 2020; Komor et al., 2016). We then annotated each sgRNA with its predicted corresponding amino acid changes and binned sgRNAs based on the most severe predicted consequence (e.g. splice site, nonsense, missense, silent; see Methods). All further screens followed this general format and annotation strategy unless otherwise noted.

**Figure 1.**
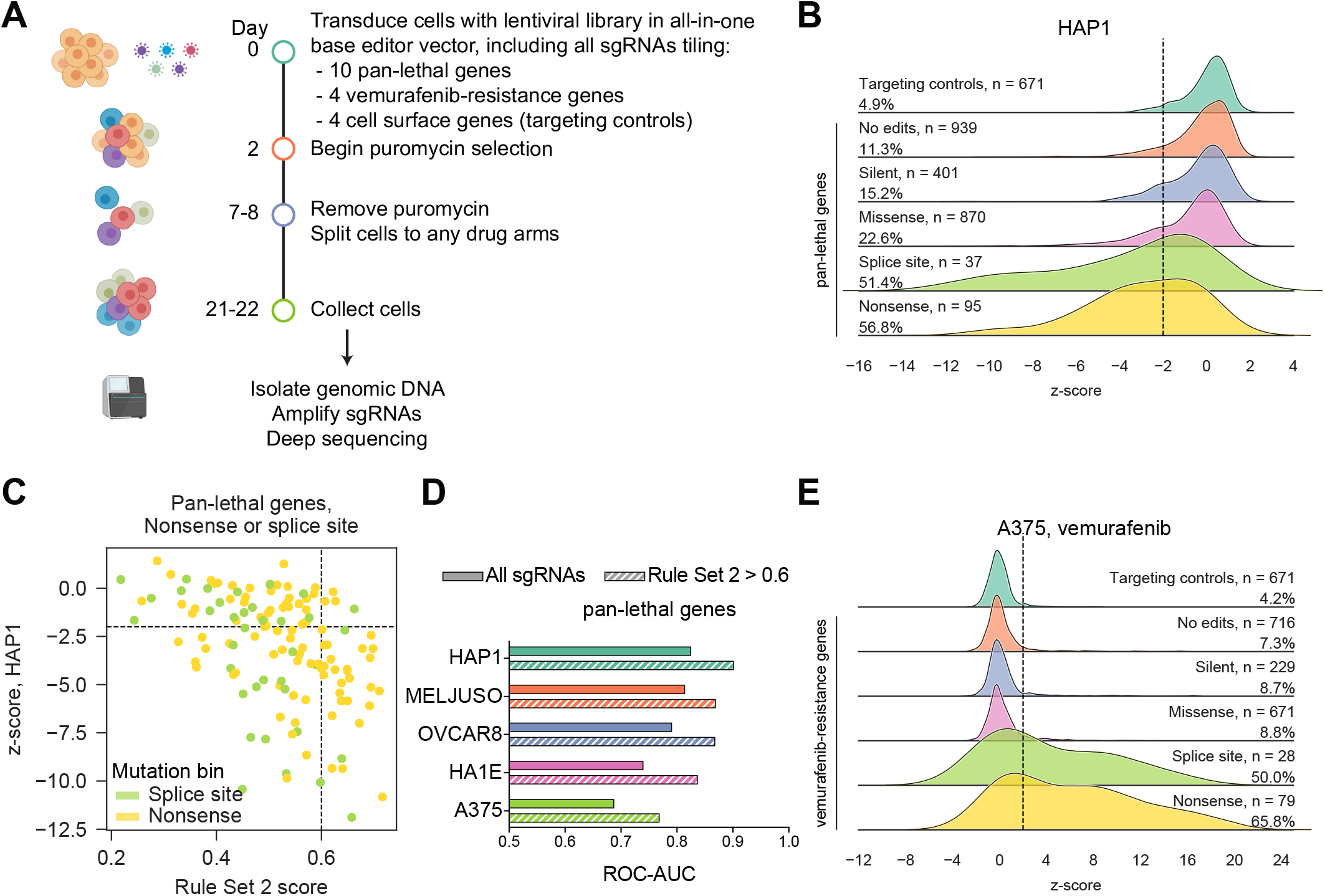
Base editors identify loss-of-function mutations in negative and positive selection screens. (A) Schematic of pooled screens with lentiviral tiling library. (B) Performance of sgRNAs targeting pan-lethal genes, grouped according to the most severe predicted mutation consequence. Targeting controls are all sgRNAs targeting non-essential cell surface markers, regardless of mutation consequence. The percentage of sgRNAs with a z-score < −2 in a given mutation bin is also reported. (C) Correlation between Rule Set 2 score and sgRNA depletion in HAP1 cells for sgRNAs predicted to introduce nonsense mutations (n = 95) or splice site mutations (n = 37) in pan-lethal genes. Dashed lines show a Rule Set 2 score of 0.6 and a z-score of −2. Pearson r = −0.44. (D) Area under the curve (AUC) values for receiver operating characteristic (ROC) curves shown in Figure S1E and S1H. (E) Performance of sgRNAs targeting NF1, NF2, CUL3, and MED12 in vemurafenib resistance screens, grouped by the most severe mutation consequence. Targeting controls are all sgRNAs targeting cell surface markers, regardless of mutation consequence. The percentage of sgRNAs with a z-score > 2 is shown.

We first asked whether sgRNAs predicted to introduce neutral or loss-of-function mutations in pan-lethal genes showed differential performance in a negative selection screen. Importantly, sgRNAs predicted to introduce no edits or only silent mutations performed similarly to targeting controls (sgRNAs targeting 4 genes encoding cell surface markers), indicating that there was not a high rate of indels or C>R editing (R = A/G) (**Figure S1B**). Encouragingly, with BE3.9max, 30.5% of sgRNAs predicted to introduce nonsense mutations and 35.1% of sgRNAs predicted to introduce splice site-disrupting mutations were depleted with a z-score < −2 (**Figure S1B**). BE4max, on the other hand, showed far less depletion of such sgRNAs, with just 9.4% and 22.2% of predicted nonsense and splice site sgRNAs scoring, respectively (**Figure S1C**). Likewise, we calculated receiver-operator characteristic (ROC) curves, defining true positives as sgRNAs targeting essential genes that introduced nonsense mutations or splice site mutations, and defining true negatives as sgRNAs targeting essential genes that introduced no edits or only silent mutations. BE3.9max outperformed BE4max by this metric (ROC-AUC = 0.69 vs. 0.62; **Figure S1D**), so we proceeded with BE3.9max for subsequent screens.

To further validate BE3.9max, we screened the same library in 4 more cell lines: MELJUSO (melanoma), OVCAR8 (ovarian cancer), HA1E (immortalized kidney), and HAP1 (near-haploid CML-derived). We again used ROC curves to quantify performance and observed similar separation between predicted neutral and loss-of-function mutations across all 5 cell lines, indicating that base editing is effective across a range of cell types of various ploidy (**Figure S1E**). In HAP1, the best-performing cell line, 56.8% of sgRNAs predicted to introduce nonsense mutations and 51.4% of sgRNAs predicted to introduce splice site mutations were depleted with a z-score < −2 (**Figure 1B**). Although a small fraction (15.2%) of sgRNAs predicted to introduce silent mutations depleted below this threshold, we expect that this is most likely due to out-of-window editing that may induce, for example, nonsense or splice site mutations (see below). Encouragingly, the performance of sgRNAs targeting pan-lethal genes was highly correlated across the 5 cell lines (Pearson r > 0.79), demonstrating consistent performance of our screens and suggesting that similar features underlie editing efficacy across cell types.

The range of phenotypic effects for sgRNAs predicted to introduce nonsense or splice site mutations in pan-lethal genes indicates that not all sgRNAs introduced base edits at a high enough frequency to be detected in a negative selection screen, so we examined several factors that could contribute to sgRNA activity. First, because we also screened the tiling library with wtCas9 in A375 and MELJUSO cells, we compared the performance of sgRNAs predicted to introduce nonsense or splice site mutations with the base editor across the two screens, as these sgRNAs would be expected to deplete with both the base editor and wtCas9. Nearly all sgRNAs depleted with z-scores < −2 in the wtCas9 condition, suggesting that the lack of a phenotype in the base editor screen was likely due to a base editor-specific effect rather than, for example, ineffective sgRNA expression or low RNA stability (**Figure S1F-G**). We also observed a correlation between the Rule Set 2 on-target score (Doench et al., 2016) and base editor performance (**Figure 1C**); filtering for high Rule Set 2 scores further improved ROC-AUC values (**Figure 1D; Figure S1H**). Filtering by predicted on-target activity may be useful for introducing a single nonsense mutation per gene (Billon et al., 2017; Kuscu et al., 2017), regardless of the position, although we note that, without base editor-specific on-target activity rules, the variability in even highly ranked sgRNAs recommends wtCas9 for standard gene knockout approaches. However, when using base editors for mutagenesis, there are typically a handful of sgRNAs (at most) that can introduce a given edit. Therefore, we opted to continue to include all possible sgRNAs in our libraries rather than pre-filter the library by predicted on-target efficacy and potentially remove active sgRNAs.

Whereas negative selection screens are useful for benchmarking the per-cell efficiency of base editing, positive selection screens are more sensitive to less frequent editing outcomes and therefore more able to detect false positives, such as base editing outside the predicted window, C>R conversions, or indels. We therefore examined the results of the positive selection screen in A375 cells for vemurafenib resistant mutations in *CUL3, MED12, NF1*, and *NF2*, and again observed a distinct separation between sgRNAs predicted to introduce no edits or silent mutations and sgRNAs predicted to introduce splice site or nonsense mutations (**Figure 1E; Figure S1I**). Importantly, fewer than 10% of sgRNAs predicted to introduce silent mutations or no edits enriched with z-score > 2, indicating that the screen had a low false positive rate. Based on these results, we conclude that base editor screens can effectively identify sgRNAs introducing loss-of-function mutations in positive and negative selection screens.

### Identification of known loss-of-function variants in *BRCA1* and *BRCA2*

Continued sequencing of cancer genomes has revealed a long tail of low-frequency mutations in tumor suppressor genes whose functional consequence, and thus clinical significance, is unclear, highlighting the need for functional characterization of variants. Loss-of-function mutations in the DNA damage repair genes *BRCA1* and *BRCA2* typically hinder cell growth and are therefore assayed by negative selection screens. We constructed 2 libraries of all possible sgRNAs tiling each gene (n = 562 sgRNAs targeting *BRCA1*, n = 589 sgRNAs targeting *BRCA2*, plus 75 non-targeting controls, 75 intergenic controls, and 32 positive controls targeting splice sites in pan-lethal genes in each library). We screened these libraries in triplicate in two cell lines (HAP1 and MELJUSO) at very high coverage (>10,000 cells per sgRNA) (**Figure 2A**). *BRCA1* and *BRCA2* are essential in HAP1 cells (Findlay et al., 2018), so cell viability (dropout) screens were conducted in these cells. MELJUSO cells were treated with either the PARP inhibitor talazoparib or the DNA-damaging agent cisplatin to sensitize cells to BRCA loss. Positive controls showed strong depletion in the untreated arms (**Figure S2A-B**) and replicates correlated strongly (Pearson r > 0.95); after averaging replicates, we also observed excellent correlation (Pearson r > 0.94) between the talazoparib and cisplatin arms in MELJUSO cells, and between the two cell lines, where the MELJUSO score was an average of the two drug conditions (Pearson r > 0.84) (**Figure S2C-D**). The consistency of sgRNA performance across treatment conditions allowed us to average sgRNA scores from each cell line, resulting in a single score per sgRNA, and compute a corresponding z-score relative to intergenic controls.

**Figure 2.**
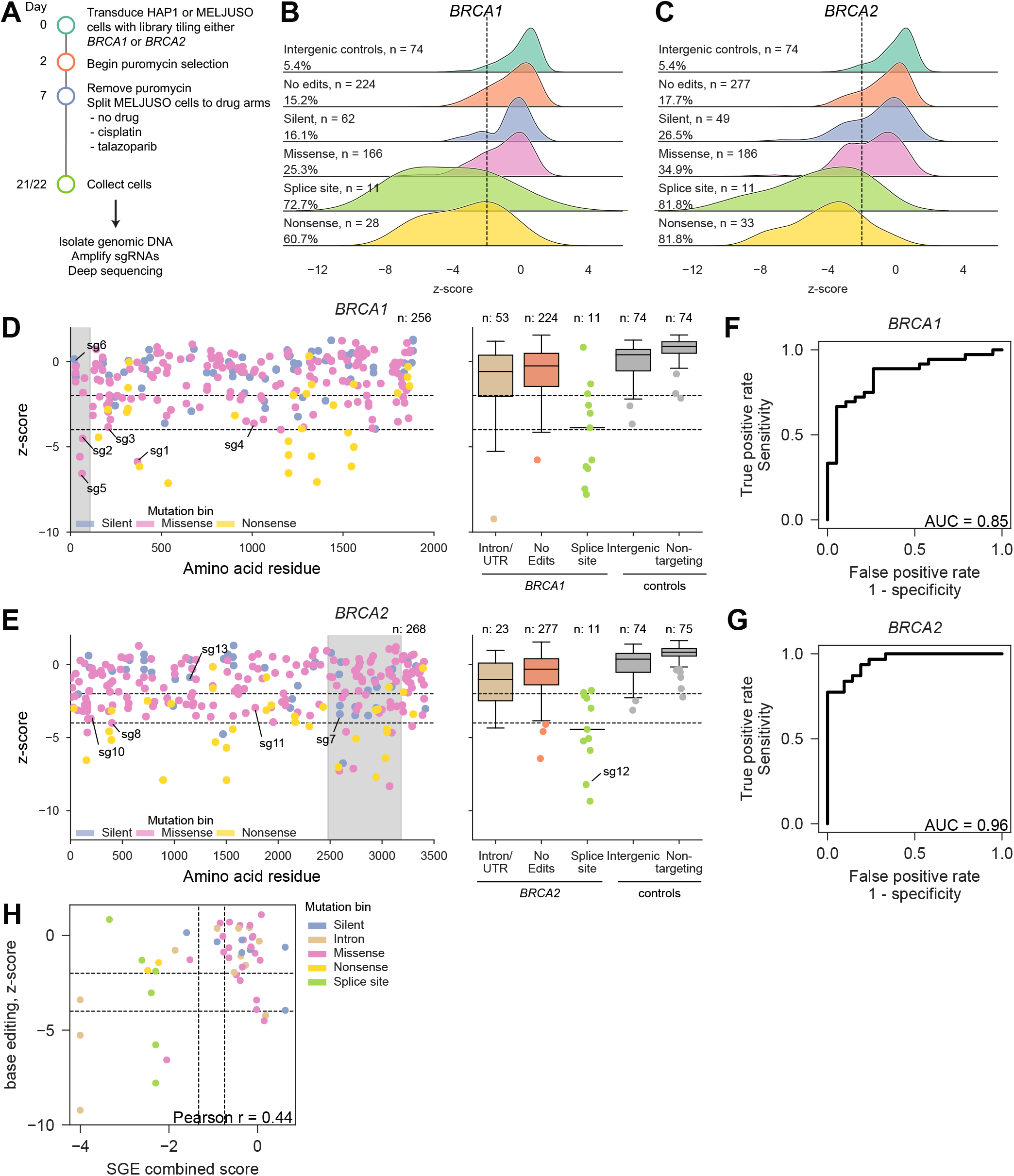
Base editor screens of BRCA1 and BRCA2 identify known loss-of-function mutations. (A) Schematic of BRCA1 and BRCA2 screens in HAP1 and MELJUSO cells. (B-C) Depletion of sgRNAs predicted to introduce nonsense mutations and splice site mutations in BRCA1 and BRCA2. Dashed line indicates the threshold between non-scoring and intermediate hit sgRNAs (z-score = −2). The percentage of sgRNAs in each category with z-score < −2 is indicated. (D-E) Performance of sgRNAs targeting BRCA1 and BRCA2, colored according to the predicted mutation bin. Dashed lines show z-score thresholds of −2 and −4; shaded areas indicate the RING domain and DNA-binding domain for BRCA1 and BRCA2, respectively. Boxes show the quartiles; whiskers show 1.5 times the interquartile range. Categories with n < 20 are shown as individual dots. (F-G) ROC curves for curated sets of pathogenic / likely pathogenic and benign / likely benign variants listed in ClinVar for BRCA1 and BRCA2. When multiple sgRNAs were predicted to introduce the same variant, the z-scores were averaged to obtain a single z-score per variant. (H) Comparison of the results of a saturation genome editing (SGE) screen of the BRCA1 RING and BRCT domains to the results of base editor screens (n = 49 sgRNAs for which all predicted edits were assayed by SGE). A combined SGE score for each sgRNA is shown and is colored by the predicted mutation bin in the base editor screen. Dashed lines show the thresholds to score in the SGE screen as described in the original publication; for the baes editor screen, dashed lines show a z-score of −2 and −4.

As before, we observed that sgRNAs predicted to introduce nonsense and splice site mutations were strongly depleted relative to sgRNAs predicted to introduce silent mutations, confirming that we could effectively assay loss-of-function mutations in *BRCA1* and *BRCA2* (**Figure 2B-C**). We binned sgRNAs by z-score, categorizing them as either strong hits (z-score < −4), intermediate hits (−4 < z-score < −2), or non-scoring (z-score > −2). 72% (44/61) and 77% (17/22) of sgRNAs predicted to introduce nonsense mutations or splice site mutations, respectively, scored as either strong or intermediate hits, in contrast to 21% (23/111) of sgRNAs predicted to introduce only silent mutations; depleted sgRNAs in the latter category likely consist of sgRNAs with out-of-window, C>R, or off-target editing. Importantly, non-scoring sgRNAs should not be conflated with benign mutations, as the primary screen is unable to rule out ineffective editing for any non-scoring sgRNA.

Of the 352 sgRNAs predicted to introduce missense mutations in *BRCA1* or *BRCA2*, 30% (107/352) scored as strong or intermediate hits. The majority of strongly-depleted missense mutations occurred in critical protein domains: in *BRCA1*, 75% (3/4) of the strong hit sgRNAs predicted to make missense mutations localized to the RING domain, which is required for BARD1 binding and E3 ubiquitin ligase activity. Likewise, in *BRCA2*, 71.4% (5/7) of the strong hit sgRNAs predicted to introduce missense mutations localized to the DNA-binding domain (**Figure 2D-E**).

We then compared our results to several orthogonal datasets. First, we curated a gold-standard set of variants that are annotated as either pathogenic / likely pathogenic (P/LP) or benign / likely benign (B/LB) in ClinVar, a database of clinically-observed genetic variation (Landrum et al., 2018), with a review status of at least one gold star (indicating a higher level of review). We filtered for variants that were predicted to be introduced by at least one sgRNA in our screen; for B/LB variants, we required that sgRNAs be predicted to introduce only the B/LB variant; otherwise, sgRNAs might deplete due to a confounding second mutation. This yielded a set of 55 variants in *BRCA1* (36 P/LP; 19 B/LB) and 52 variants in *BRCA2* (31 P/LP; 21 B/LB). When variants were introduced by multiple sgRNAs, we averaged the scores to obtain a single score for each variant. We found that base editor screens distinguished P/LP from B/LB variants with an AUC of 0.85 for *BRCA1* and 0.96 for *BRCA2* (**Figure 2F-G**). At a z-score threshold of −2, the *BRCA1* screen had a sensitivity of 0.70 and a specificity of 0.84; the *BRCA2* screen had a sensitivity of 0.84 and a specificity of 0.86. These findings indicate that base editor screens can identify mutations of functional importance with high precision.

For *BRCA1*, we also compared our results to existing data from a saturation genome editing (SGE) screen of single nucleotide variants in the 13 exons encoding the RING and BRCT domains (Findlay et al., 2018). We calculated a combined SGE score for all sgRNAs that were predicted to introduce mutations also assayed in the SGE dataset; to calculate a combined score, we obtained the SGE scores for all the mutations predicted to be made by a given sgRNA, then took the minimum (reasoning that, in most cases, the mutation with the most deleterious effect should drive the behavior of the sgRNA). We excluded any sgRNAs from this comparison that were predicted to make mutations not assayed by SGE. Overall, we calculated a combined SGE score for 51 sgRNAs and observed a modest correlation between the two datasets (Pearson r = 0.44; **Figure 2H**). We binned SGE scores into loss-of-function, intermediate, and functional based on the definitions in the original publication, and observed that 56.8% of sgRNAs (29/51) either scored in both datasets or neither dataset. Very few sgRNAs (2/51) scored strongly in the base editor screen but not at all in the SGE screen, consistent with the low false positive rate observed in previous experiments.

### Validation of *BRCA1* and *BRCA2* loss-of-function alleles

Despite good agreement with gold-standard datasets, it is difficult to draw definitive conclusions about individual mutations based on the results of the primary screen alone because sgRNAs may deplete due to reproducible but unanticipated effects, such as out-of-window editing, C>R editing, indels, or editing at off-target sites. Therefore, we individually validated 13 sgRNAs (sg1-sg13). Since many nonsense mutations in *BRCA1* and *BRCA2* have been well characterized, we focused on sgRNAs predicted to introduce missense or silent mutations. In order to determine the dynamic range of our assay, we selected sgRNAs with a range of activity in the primary screen. We also selected several sgRNAs of interest, such as 1 of the 2 sgRNAs that scored in the base editor screen but not the SGE screen (sg2) and a strong hit sgRNA targeting outside the RING domain of *BRCA1* (sg1); additionally, we included an empty vector control (NTC-1) that did not contain an sgRNA sequence. We individually cloned these sgRNAs into the same all-in-one BE3.9max lentiviral vector used for screens, transduced HAP1 cells with each vector in duplicate, selected for transduced cells with puromycin, and cultured cells for a total of 3 weeks to allow alleles to enrich or deplete, collecting cells at 7, 14, and 21 days post-transduction (**Figure 3A**). We then used custom primers for each sgRNA to amplify the genomic region surrounding the edit site and subjected the amplicons to deep sequencing.

**Figure 3.**
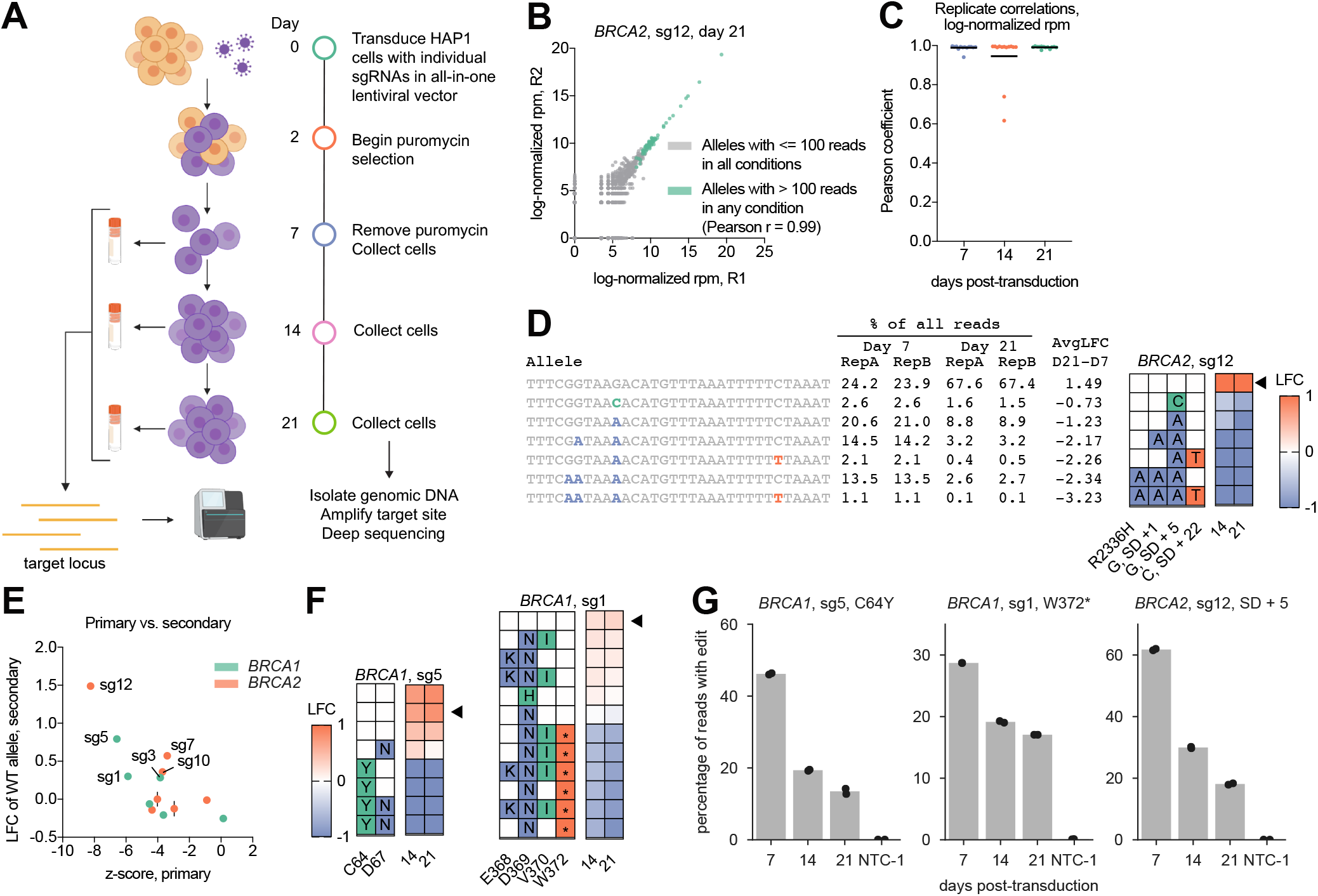
Validation of 13 sgRNAs targeting BRCA1 and BRCA2. (A) Schematic of validation experiments in HAP1 cells. (B) Replicate correlation for allele-level log2-normalized reads per million for sg12 at 21 days post-transduction. Pearson correlation between alleles with at least 100 reads in any sg12-treated condition is reported. (C) Distribution of Pearson coefficients for allele-level replicate correlations for sg1-sg13 at 7, 14, and 21 days post-transduction. The mean is shown in black. (D) Abundance and log-fold change (LFC) of alleles for sg12-treated cells at 7 or 21 days post-transduction. Only alleles with at least 1% abundance in at least one condition are included on the heatmap. The amino acid and splice site edits are summarized in the map on the right; average LFC for days 14 and 21 (relative to day 7) is shown in the heatmap. The wild-type (unedited) allele is indicated with a black triangle. “SD” indicates splice donor. (E) Comparison between sgRNA performance in the primary screen (z-score) and secondary validation (LFC of the wild-type allele at 21 days post-transduction). Error bars show the range of 2 biological replicates in the secondary validation. Guides with a positive LFC of the wild-type allele are labeled. (F) Log-fold change at days 14 and 21 (relative to day 7) of alleles in sg5- and sg1-treated cells. (G) Percentage of all sequencing reads containing the indicated mutation at each timepoint. Dots show n = 2 biological replicates. Reads containing indels were removed from consideration. “SD” indicates splice donor. NTC-1 indicates a non-targeting (no sgRNA) control.

We first assessed base editing efficiency across the 13 sgRNAs tested (using the early, day 7 timepoint to mitigate the effect of sgRNA dropout). For each sgRNA, we calculated the editing efficiency for each C in a broad window around the sgRNA (positions −10 to 20, where position 1 is the first nucleotide of the protospacer and positions 21 - 23 are the PAM sequence). We observed that the strongest editing was C>T editing in the target window (positions 4 to 8; median = 46.6% C>T editing), with low levels of C>R editing and deletions (**Figure S2E**). C>T editing outside the canonical window occurred at a lower but detectable rate. Notably, the 13 sgRNAs selected for validation may be enriched for active sgRNAs relative to the library overall, which could inflate the observed efficiency of base editing. Still, the overall trends of editing are consistent with previous reports and provide confidence that long-term expression of lentiviral-delivered base editors do not induce widespread indels or C>R editing.

Next, we sought to leverage the allele-level information produced by next-generation sequencing to separate causal and passenger mutations for each sgRNA. We filtered out alleles with <100 reads in all samples and observed strong correlation of log-normalized allele read counts between replicates (**Figure 3B-C**), indicating that patterns of editing were highly reproducible. We then calculated a log-fold change for each allele by comparing the allele frequency at days 14 and 21 to the allele frequency at day 7 (**Figure 3D**) and averaged replicates to obtain an average log-fold change for each allele. We observed the expected negative correlation between the primary screening results and the log-fold change of wildtype alleles in the secondary validation (Spearman r = −0.53; **Figure 3E**): if an sgRNA scored with a negative log-fold change in the primary screen, then in the secondary validation, the wildtype allele should out-compete the deleterious edit and therefore enrich with a positive log-fold change. Overall, 8/13 sgRNAs replicated the results of the primary screen by this metric; 6/8 scored as hits in both experiments, whereas 2/8 did not score in either. The remaining 5/13 sgRNAs, including the guide that scored in the base editor screen but not the SGE screen (sg2), represent false positives of the primary screen. This small sample is likely not representative of the overall false positive rate, however, as we chose to validate a non-random sample of sgRNAs, including several intermediate hits from the primary screen.

To determine the mutations driving the phenotype of the 6 sgRNAs that scored in both the primary and secondary experiments, we visualized the amino acid edits in each allele alongside the log-fold change (**Figure 3D**). The three sgRNAs with the strongest phenotype in both the primary and secondary screens are highlighted here. In *BRCA1*, sg5 scored as a strong hit in the screen (z-score = −6.57), and all of the alleles with a negative log-fold change introduced the predicted C64Y mutation (rs55851803; **Figure 3F**), which is classified as Pathogenic in ClinVar and has been shown to disrupt E3 ubiquitin ligase activity (Ruffner et al., 2001). This edit decreased from 46.2% of reads on day 7 post-transduction to 13.5% on day 21 (**Figure 3G**). Importantly, this example highlights that an initial editing efficiency of ~50% at day 7 was sufficient to see substantial depletion in the primary screen. The second strongest validation sgRNA targeting *BRCA1*, sg1, was predicted to introduce D369N and V3701 mutations; however, in the validation experiments, this guide showed distinct depletion of alleles containing a nonsense mutation at W372 (introduced by C>T edits at position −2), highlighting the importance of validating actual edits rather than relying on predictive heuristics alone (**Figure 3F-G**).

Similarly, in *BRCA2*, the most depleted sgRNA in the validation (sg12) showed depletion of alleles containing a G>A mutation at the canonical splice donor site of exon 13 (rs397507891), which is listed as “Pathogenic” in ClinVar (**Figure 3D; Figure 3G**). Interestingly, this sgRNA also gave rise to a second depleted allele that contained an intact splice donor site but included a G>A mutation 5 nucleotides into the intron (“SD + 5”, where “SD” denotes a splice donor site; rs81002816), which is considered to have “Conflicting interpretations of pathogenicity” in ClinVar. An analysis of splicing intolerance using ExAC data found that the SD + 5 site was significantly intolerant of non-G nucleotides (Zhang et al., 2018). Although we cannot conclusively determine the functional consequence of this mutation without further validation, our data are consistent with the conclusion that G>A mutations at this SD + 5 site of *BRCA2* exon 13 are loss-of-function and disrupt splicing.

### Mutagenesis of *MCL1* and *BCL2L1* in the presence of targeted inhibitors

We next sought to use base editor screens to probe drug-target interactions, hypothesizing that mutagenesis of drug targets could help identify specific mutations that confer drug sensitivity or resistance. Detailed understanding of such mutations would be useful for stratifying patient populations and could inform rational design of future drugs. *MCL1* and *BCL2L1*, a pair of anti-apoptotic genes often upregulated in cancer, share a synthetic lethal relationship (Han et al., 2017; Najm et al., 2017) and each have targeted inhibitors (S63845, hereafter “MCL1-i”; A-1331852, hereafter “BCL2L1-i”) (Kotschy et al., 2016; Leverson et al., 2015). Thus, inhibition of one target protein allows for screens in which loss-of-function mutations in the other gene will deplete. Additionally, we reasoned that we could identify mutations that confer resistance to either inhibitor while still depleting loss-of-function variants by treating cells with intermediate doses of both inhibitors (**Figure 4A**).

**Figure 4.**
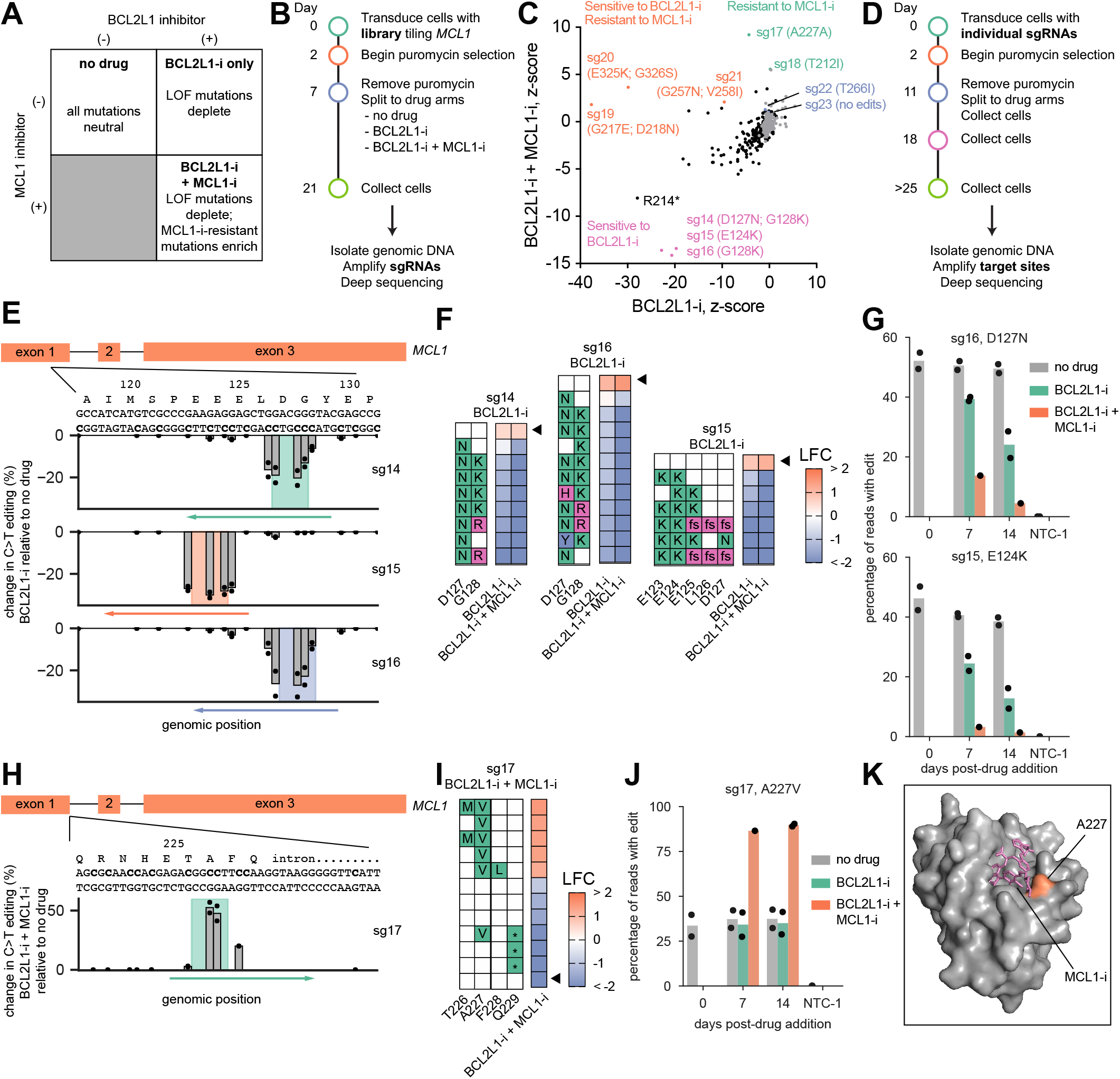
Base editor screens uncover missense mutations in MCL1 conferring sensitivity and resistance to BH3 mimetics. (A) The synthetic lethal relationship between MCL1 and BCL2L1 dictates the expected behavior of MCL1 mutations in each condition of the screen. LOF = loss-of-function. (B) Timeline of pooled screen. (C) Performance of sgRNAs in the presence of BCL2L1-i (A-1331852) alone versus BCL2L1-i and MCL1-i (S63845). Guides selected for validation are labeled with their predicted mutations and colored by their performance in the primary screen. Negative control sgRNAs (non-targeting and intergenic) are shown in gray. (D) Timeline of validation of individual sgRNAs. (E) Validation of sg14, sg15, and sg16. Schematic of sgRNA targets in MCL1 (above) and difference in the percentage of C>T editing in BCL2L1-i treated cells relative to untreated cells. Shaded boxes show the window of expected editing for each sgRNA; arrows show the location of each sgRNA, with the arrowhead at the PAM-proximal end of the protospacer. Dots show n = 2 biological replicates. (F) Identification of likely-causal alleles from deep sequencing data. For each allele, the log-fold change (LFC) is shown for drug-treated cells relative to untreated cells, and the triangle indicates the wildtype (unedited) allele. Amino acid changes in each allele are depicted on the left; “fs” = frameshift. Only alleles with >1% of all reads in at least one condition were included on the heatmap. (G) Percentage of reads containing D127N (above) and E124K (below) at each of 3 timepoints and 3 drug conditions. Dots show n = 2 biological replicates. The BCL2L1-i and MCL1-i co-treatment condition has only one replicate, due to low cell numbers. NTC-1 indicates an empty vector control. (H) Validation of sg17. Schematic of sg17 targeting MCL1 (above) and difference in the percentage of C>T editing in BCL2L1-i and MCL1-i co-treated cells relative to untreated cells. Shading shows the window of expected editing. (I) Allele analysis reveals A227V as the likely causal mutation, as in (F). (J) Fraction of reads containing A227V at each of 3 timepoints and 3 drug conditions. Dots show n = 2 biological replicates. The BCL2L1-i + MCL1-i sample at 7 days post-drug addition has only one replicate, due to low cell numbers. Reads containing indels were not considered to be edited. NTC-1 indicates an empty vector control. (K) A227 (orange) directly contacts S63845 (“MCL1-i”, pink) in the structure of MCL1 bound to S63845 (PDB: 5LOF).

To explore this approach, we constructed a library of 210 sgRNAs tiling *MCL1*, as well as 150 negative controls (non-targeting and intergenic) and 32 positive controls targeting splice sites in pan-lethal genes. We conducted screens in duplicate at very high coverage (>10,000 cells per sgRNA) in MELJUSO cells across 3 conditions: no drug, BCL2L1-i alone at a dose of 250 nM, or co-treatment with MCL1-i and BCL2L1-i, each at a dose of 62.5 nM (**Figure 4B**). We observed excellent correlation between untreated replicates (Pearson r = 0.99) and substantial depletion of pan-lethal splice site controls, indicating that the technical quality of the screen was high (**Figure S3A**). We then compared the performance of sgRNAs in our drug treatment arms (BCL2L1-i alone versus co-treatment with BCL2L1-i and MCL1-i, **Figure 4C**). Two sgRNAs (sg17, sg18) conferred strong resistance (z-score > 4) to cells treated with both inhibitors; each of these sgRNAs is predicted to introduce missense mutations in the BH3 helix. Several sgRNAs sensitized cells to BCL2L1-i treatment, with or without the addition of MCL1-i. These sgRNAs include an sgRNA predicted to introduce a nonsense mutation at R214, and 3 sgRNAs (sg14, sg15, sg16) predicted to introduce missense mutations in residues 124-128 in the PEST domain of MCL1. Unexpectedly, several sgRNAs (sg19, sg20, sg21) strongly sensitized cells to BCL2L1-i treatment, but this effect was lost upon co-treatment with MCL1-i, suggesting that the editing products of these sgRNAs could affect previously unappreciated residues for MCL1 function.

To identify the mutations driving these phenotypes, we selected 10 sgRNAs to validate, including 8 sgRNAs that showed a strong phenotype in one or both of the drug arms (sg14-21) and 2 sgRNAs (sg22, sg23) that did not score strongly in either drug condition; we also included a negative control sgRNA targeting EGFP (NTC-2). As before, we transduced cells with these sgRNAs individually, cultured cells in the presence of either single or dual inhibitors for at least 2 weeks, and sequenced the target loci (**Figure 4D**). Across all 10 validated sgRNAs in *MCL1*, the log-fold change of the WT allele in the validation experiment negatively correlated with the z-score of the sgRNA in the primary screen (**Figure S3B-C**) and replicates were generally well correlated at the allele level (**Figure S3D**).

The 3 sgRNAs targeting the PEST domain (sg14-16) introduced several distinct deleterious edits. Editing with sg15 led to multiple distinct genotypes with E>K mutations introduced across residues 123, 124, and 125, whereas editing with sg14 or sg16 introduced D127N and G128K/R mutations. The D127N mutation arising from sg14 and sg16 was sufficient to strongly sensitize cells to BCL2L1-i treatment, suggesting that it confers a loss-of-function phenotype (**Figure 4E-F**). Mutations introduced by sg14-16 depleted dramatically in both the BCL2L1-i-treated and BCL2L1-i and MCL1-i-treated arms; for example, with sg16, the D127N mutation depleted from 52.1% of reads prior to drug treatment to 4.5% after 14 days of combination drug treatment (**Figure 4G**). D127 is known to be the site of caspase cleavage (Clohessy and Zhuang, 2004; Herrant et al., 2004), and the “EELD” sequence at residues 124-127, which is partially disrupted by all 3 sgRNAs, has been shown to be required for Tom70-dependent mitochondrial targeting (Chou et al., 2006), suggesting that these point mutations may disrupt MCL1 function by altering either the cleavage or localization of MCL1.

We next examined the samples treated with both BCL2L1-i and MCL1-i. The top hit from the primary screen, sg17, showed clear enrichment of an A227V mutation (**Figure 4H-I**); reads with these edits enriched from 33.6% abundance prior to drug treatment to 89.7% after 14 days of combination treatment, with no similar effect in the cells treated with BCL2L1-i alone (**Figure 4J**). This residue lies in the MCL1-i binding pocket and directly contacts the small molecule, suggesting that this mutation confers resistance by directly disrupting binding to MCL1 (**Figure 4K**). Guides 19 and 21 also induced resistance through mutations adjacent to the MCL1-i binding site at G217/D218 and G257, respectively (**Figure S4A-F**). Interestingly, mutations at G217/D218, in the BH3 helix, and D256, in the BH1 helix, sensitized cells to BCL2L1-i treatment alone. The similarity of this cluster of mutations that confer resistance to MCL1-i and BCL2L1-i suggests that these are key residues for MCL1 function. We also uncovered several mutations outside the MCL1-i binding site that conferred resistance, including T212I (sg18) and an allele with several mutations near the C-terminus (E322K, E325K, G326N; sg20) (**Figure S4G-L**). Notably, the E325K mutation alone sensitized cells to BCL2L1-i, suggesting that this residue in the C-terminus of MCL1 may be a linchpin in the functional role of this domain. In sum, through a small screen and focused validation, we uncovered numerous single, double, and triple mutants in *MCL1*, including both loss-of-function and drug-resistant mutants, highlighting the utility of these screens for biological discovery.

We conducted an analogous tiling screen in *BCL2L1* (**Figure 5A; Figure S5A**), treating cells with either MCL1-i alone or both MCL1-i and BCL2L1-i at an intermediate dose. Again, we identified a number of sgRNAs that depleted very strongly in the single inhibitor arm, including all 13 sgRNAs predicted to introduce nonsense or splice site mutations (**Figure 5B**). We additionally identified 8 sgRNAs that enriched highly (z-score > 4) in the MCL1-i and BCL2L1-i treated arm; interestingly, the top 5 sgRNAs were all predicted to make no edits or silent mutations, suggesting the presence of out-of-window or C>R editing (**Figure 5C**). We therefore validated 5 sgRNAs that conferred resistance to co-treatment (sg24-28; **Figure 5C-D**). Replicates correlated well (**Figure S5B**) and we observed a negative correlation between performance of sgRNAs in the primary screen and log-fold change of the WT allele in the validation, indicating that the validation recapitulated the results of the primary screen (Spearman r = −0.70; **Figure S5C**).

**Figure 5.**
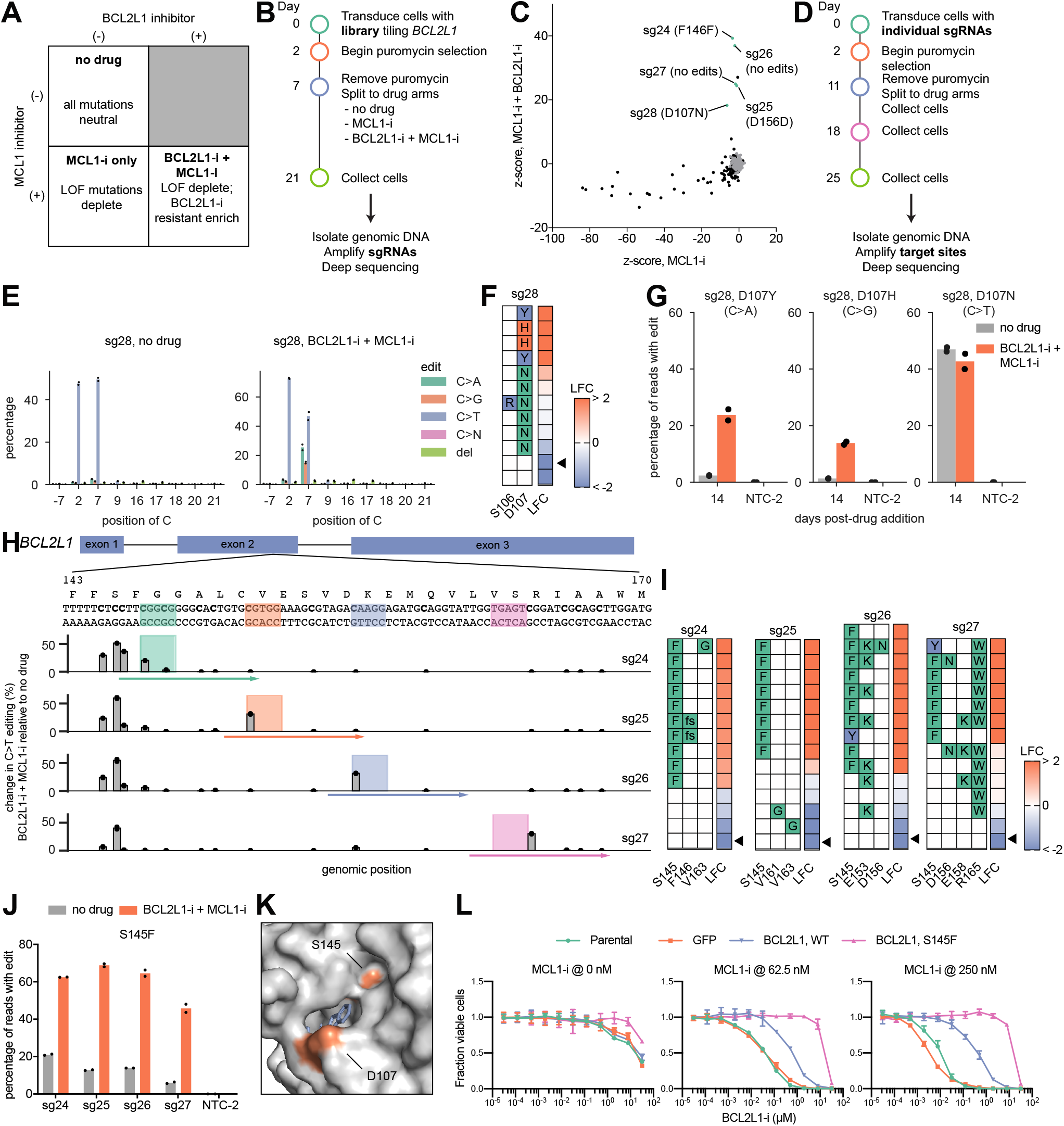
Mutagenesis of BCL2L1 reveals drug resistant point mutations. (A) The synthetic lethal relationship between MCL1 and BCL2L1 dictates the expected behavior of BCL2L1 mutations in each condition of the screen. LOF = loss-of-function. (B) Timeline of pooled screen. (C) Performance of sgRNAs in the presence of MCL1-i treatment versus BCL2L1-i and MCL1-i co-treatment. Guides selected for validation are colored and labeled. Negative controls (non-targeting and intergenic) are shown in gray. (D) Timeline of validation of individual sgRNAs. (E) Comparison of the types of edits generated by sg28 in the presence or absence of drug treatment. Dots show n = 2 biological replicates. (F) Identification of likely-causal alleles from deep sequencing data. For each allele, the log-fold change (LFC) is shown for drug-treated cells relative to untreated cells, and the triangle indicates the wildtype (unedited) allele. Only alleles with >1% of all reads in at least one condition were included on the heatmap. (G) Percentage of reads containing the indicated edits in the presence and absence of drug treatment. Reads containing indels were not considered to be edited. Dots show n = 2 replicates. NTC-2 indicates time-matched non-targeting control (EGFP-targeting sgRNA). (H) Deep sequencing of the genomic region surrounding the edit sites for sgRNAs 24 - 27 reveals that alleles harboring a S145F mutation enrich in the combination-treated condition. Values plotted are the average of two replicates. Shaded regions indicated the expected editing window. (I) Identification of likely causal alleles, as in (F). (J) Percentage of reads containing the indicated edit in the presence and absence of drug treatment. Dots show n = 2 replicates. Reads containing indels were not considered to be edited. NTC-2 indicates time-matched non-targeting control (EGFP-targeting sgRNA). (K) Location of S145 in the crystal structure of BCL2L1 bound to A-1155463, a compound that is structurally similar to A-1331852 (the BCL2L1 inhibitor used in experiments) (PDB: 4QVX). (L) Overexpression of a BCL2L1 ORF with the S145F mutation shifts the dose-response curve of BCL2L1-i, shown at 3 concentrations of MCL1-i. Cell Titer Glo measurements were taken after 4 days of treatment and are normalized to cells not treated with BCL2L1-i. Error bars represent the standard deviation of n = 4 technical replicates.

The validation of these 5 sgRNAs underscores the importance of directly sequencing the target site prior to drawing conclusions about the mutations responsible for the phenotype. In the untreated condition, sg28 introduced the expected edit: C7 of the sgRNA, corresponding to a D107N mutation, was found in nearly 50% of reads, with only ~5% of reads containing C>A edits, C>G edits, or deletions at the C7 position. Upon BCL2L1-i and MCL1-i co-treatment, however, the C>A and C>G edits enriched strongly, to 25% and 15% of all reads, respectively (**Figure 5E**). Thus, although sg28 did not introduce a high fraction of C>R editing in the untreated arm, these alleles enriched under strong positive selection. Once validated, however, the heterogeneity produced by this sgRNA proved useful, revealing two resistance mutations: D107H and D107Y (**Figure 5F-G**). The structure of BCL2L1 bound to A-1155463 (Tao et al., 2014), a BCL2L1 inhibitor that is structurally similar to the BCL2L1-i we used (A-1331852), reveals that D107 lies in the drug binding pocket of BCL2L1 (**Figure 5K**), where substitution with a bulkier amino acid (H, Y) may confer stronger resistance than a more compact one (N, D).

We next examined the additional 4 sgRNAs targeting *BCL2L1*, sg24-27, all of which target a similar region of *BCL2L1* (**Figure 5H**). These sgRNAs displayed a distinct pattern of editing: in the untreated condition, each sgRNA showed not only the predicted editing in or near the expected window but also a detectable level of C>T editing at two particular C’s upstream of the sgRNA. This editing ranged from 26.1% of reads for sg24 (for which the edits correspond to positions −2 and 0 relative to the sgRNA, i.e. 4-6 nucleotides upstream of the predicted window) to 6.2% of reads for sg27 (for which the edits correspond to positions −52 and −50, i.e. 54-56 nucleotides upstream of the predicted window). Editing this far outside the window was highly unusual among the sgRNAs we validated, and may be due to locus-specific base editing patterns; for example, the sequence immediately surrounding these nucleotides is T-rich, and a TC motif has been documented as a preferred substrate for rAPOBEC1 (Beale et al., 2004). In a negative selection screen, editing at a low frequency would likely make little impact on the results; however, in this strong positive selection screen, these edits enriched in the co-treatment condition (**Figure 5H**). Alleles with mutations at these extremely PAM distal positions enriched if and only if they also contained a mutation at S145, most commonly S145F (**Figure 5I-J**). As with D107, S145 lies directly in the binding pocket (**Figure 5K**). We confirmed this result by overexpressing an ORF containing the S145F mutation, and observed a substantial shift in the dose-response curve to BCL2L1-i in the presence of MCL1-i (**Figure 5L**). These two results underscore the importance of determining the actual edit introduced by a base editor in positive selection screens, because unpredictable, low-frequency edits can drive the resulting phenotype. However, with proper validation, these sgRNAs identified functionally-relevant point mutations in *BCL2L1*.

### Mutagenesis of *PARP1*

We then asked whether a similar approach could uncover differences between multiple drugs targeting the same protein. We designed a library including all possible sgRNAs (n = 656) tiling *PARP1*, an important oncology target due to observed synthetic lethality in tumors with loss-of-function mutations in *BRCA1* and other DNA damage genes (Farmer et al., 2005; Zimmermann et al., 2018). We screened this library in HAP1 cells with 5 clinical PARP inhibitors (PARPi): niraparib, olaparib, rucaparib, talazoparib, and veliparib (**Figure 6A; Figure S6A**). As anticipated (Pettitt et al., 2018), we observed that sgRNAs predicted to introduce nonsense or splice site mutations in *PARP1* conferred resistance to all PARP inhibitors (**Figure S6B**). To identify residues that behaved differently across the inhibitors, we focused on sgRNAs that conferred resistance (z-score > 2) to at least one drug and sensitivity (z-score < −2) to another, and identified 10 sgRNAs that met these criteria (**Figure 6B**). Interestingly, 4 of these sgRNAs targeted the same residue, G646, which falls just before the catalytic domains of PARP1. We also identified 11 sgRNAs that conferred sensitivity to all 5 PARP inhibitors (z-score < −2); of these sgRNAs, 3 were predicted to edit the A774 residue and 3 were predicted to edit at or near the D692 residue, which also falls near the splice junction at the beginning of exon 15. Overall, the set of sgRNAs that sensitized to all inhibitors was enriched (82%, 9/11) for guides targeting the catalytic domains of PARP1 (residues 662-1014) (**Figure 6B; Figure S6C**).

**Figure 6.**
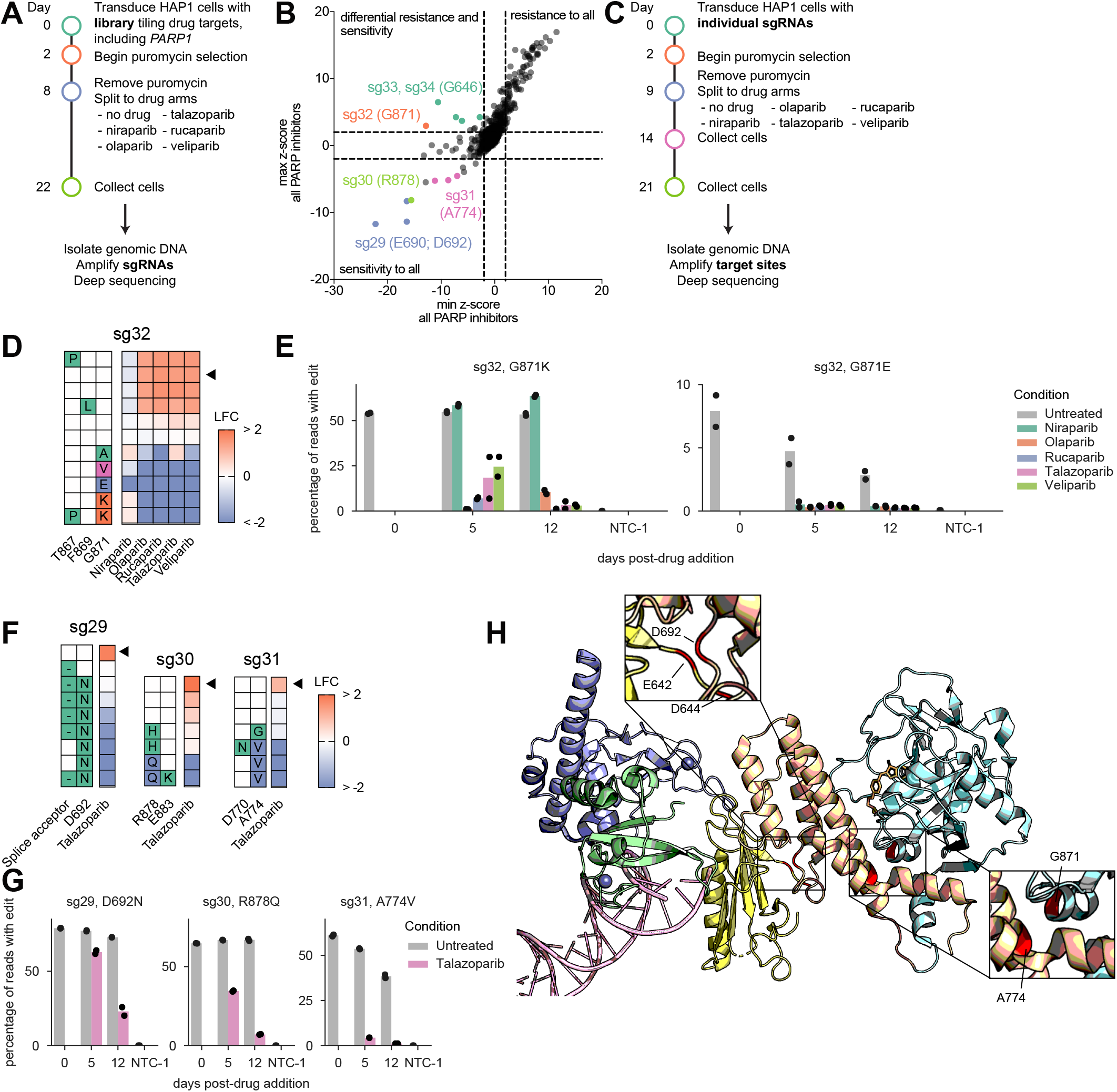
Mutagenesis of PARP1 reveals missense mutations in the catalytic domain that sensitize to PARP inhibition. (A) Timeline of PARPi screens in HAP1 cells. (B) Performance of sgRNAs targeting PARP1 in the primary screen across all 5 PARPi conditions (minimum z-score of all 5 PARPi versus maximum z-score of all 5 PARPi). A subset of the highlighted sgRNAs, displaying either sensitivity to all PARPi or differential sensitivity to PARPi, were selected for individual validation. (C) Timeline of validation of individual sgRNAs with PARPi. Guides that conferred differential sensitivity in the primary screen were validated with all 5 PARPi; guides that sensitized to all PARPi in the primary screen were validated with talazoparib only. (D) Alleles present in > 1% of reads in at least one replicate of sg32-treated cells. Amino acid changes for each allele are shown on the left; the average log-fold change of each allele (comparing drug-treated to untreated cells at 12 days post-drug addition) is shown on the right. The wildtype (unedited) allele is marked with a black triangle. (E) Percentage of all sequencing reads containing the indicated edit for each timepoint and drug condition in sg32-treated cells. Reads containing indels were not considered to be edited. NTC-1 indicates an untreated, non-targeting control (no sgRNA), sequenced at 12 days post-drug addition. (F) Alleles present in > 1% of reads in at least one replicate of sg29-, sg30-, and sg31-treated cells. Amino acid and intronic changes for each allele are shown on the left; the average log-fold change of each allele (comparing drug-treated to untreated cells at 12 days post-drug addition) is shown on the right. The wildtype (unedited) allele is marked with a black triangle. (G) Percentage of all sequencing reads containing the indicated edit for each timepoint and drug condition. Reads containing indels were not considered to be edited. NTC-1 indicates an untreated, non-targeting control (no sgRNA), sequenced at 12 days post-drug addition. (H) Validated mutations in PARP1 are shown in red on the crystal structure of PARP1 bound to a single-stranded break (PDB: 4OQB). From left to right: Zn2 domain (blue), Zn3 domain (green), DNA (pink), tryptophan-glycine-arginine (WGR) domain (yellow), helical domain (HD; tan), ADP-ribosyl transferase (ART) domain (light blue). Top left inset shows the interface of the HD and WGR domains; bottom right inset shows the αF/αJ interface of the HD and ART domains.

Next, to identify the underlying mutations causing PARPi sensitivity or differential response, we selected 8 sgRNAs to validate individually, including 3 sgRNAs that conferred sensitivity to PARPi, 3 sgRNAs that conferred differential response to PARPi, and 2 controls predicted to introduce either a nonsense mutation or a silent mutation in PARP1 (sg29-36). As before, we transduced cells with each of these individual sgRNAs, selected with puromycin, and cultured cells for 2 weeks in the presence of PARPi. For sgRNAs that conferred sensitivity to all PARP inhibitors, we divided cells into an untreated and a talazoparib-treated condition; for each of the sgRNAs that conferred differential response to the different PARP inhibitors, we repeated each of the 6 arms from the primary screen. We isolated genomic DNA at 3 time points (pre-treatment, 5 days of treatment, and 12 days of treatment) and deeply sequenced the target loci to determine the edits for each treated and untreated condition for each sgRNA.

As before, we observed strong allele-level replicate correlation (**Figure S6D**) and the expected negative correlation between z-score in the primary screen and log-fold change of the WT allele in the validation (**Figure S6E**; Spearman r = −0.98). Additionally, nonsense mutations in the positive control conferred resistance to talazoparib (**Figure S6F-G**). Examination of the individual alleles revealed numerous missense mutations, spanning 3 domains of PARP1, that sensitize to PARPi. The 2 sgRNAs targeting near G646 showed a distinct pattern: double mutant alleles (D644N and G646N/S) depleted strongly in the rucaparib and veliparib conditions, whereas alleles containing an E642K mutation enriched slightly in the talazoparib and niraparib arms (**Figure S6H-I**). Guide 32, which was enriched in the niraparib arm of the screen but depleted in all others, introduced edits mainly at G871. G871K mutants mirrored the behavior of the primary screen and showed slight resistance to niraparib only, while mutation to an acidic residue (G871E) conferred sensitivity to all 5 inhibitors (**Figure 6D-E**). Likewise, R878Q (sg30), A774V (sg31), and D692N (sg29) mutations all strongly sensitized cells to talazoparib (**Figure 6F-G**).

Visualization of these residues in the existing crystal structure of PARP1 bound to a single-stranded DNA break reveals that many of these mutations occur near the interdomain interfaces, which are thought to be critical for PARP’s response to DNA damage (Zandarashvili et al., 2020). For instance, A774 and G871 interact to form the αF/αJ interface (**Figure 6H**), one of two main interfaces between the HD and ART domains of PARP1 (Dawicki-McKenna et al., 2015). Point mutations at these residues (A774S/L; G871S/L) have been shown to disrupt the contacts between the HD and ART domains, leading to PARP1 activation (Dawicki-McKenna et al., 2015). E642, D644, and D692 also fall near the interface of the WGR and HD domains (**Figure 6H**). Interestingly, another point mutation (R591C) at the interface between the WGR domain and the HD domain has been clinically observed and was shown to disrupt PARP trapping, causing acquired olaparib resistance (Pettitt et al., 2018). Our finding that nearby point mutations can have the opposite effect - sensitization to PARPi - underscores the complex role of interdomain contact in PARPi.

In sum, in modifier screens tiling genes that encode drug targets with and without corresponding inhibitors, we demonstrate that base editor screens effectively identify point mutations that modulate drug response. Importantly, unlike screens that rely on pseudo-random mutagenesis (Chen et al., 2020; Donovan et al., 2017; Hess et al., 2016; Neggers et al., 2018), base editor technology is efficient enough to allow for negative selection screens, enabling the identification of mutations that sensitize cells to drug treatments, which may be particularly useful for guiding the design of future small molecule inhibitors.

### Functional screens of 52,034 variants in ClinVar

The most common category of SNPs in ClinVar are variants of unknown significance (VUS), emphasizing the need for massively parallel strategies to associate genotype to phenotype. Of the 388,496 single nucleotide variants in the database at the time of library design, we identified 52,034 that could be introduced with C>T base editing technology across 3,584 genes (i.e. C>T or G>A variants that fell within the canonical editing window for at least one sgRNA; **Figure 7A**). We designed a library of 68,526 sgRNAs predicted to introduce these variants, including negative controls (non-targeting, intergenic, and splice site-targeting in non-essential genes) and positive controls (splice site-targeting in essential genes). To account for bystander mutations, we used Ensembl’s variant effect predictor (VEP) (McLaren et al., 2016) to annotate each sgRNA with all predicted edits and classified each guide by the most severe predicted impact, including both the intended edit listed in ClinVar and any bystander mutations predicted to occur in the base editing window, only annotating edits in a non-GC motif (due to low editing efficiency in GC motifs). We then screened this library in triplicate in HT29 (colorectal adenocarcinoma) and MELJUSO (melanoma) cells (**Figure S7A**). We included screening arms with low-doses of cisplatin and hygromycin (**Figure 7B**), to trigger DNA damage and translational stress, respectively, reasoning that the viability requirements of many genes may only be revealed under more challenging growth conditions, as has been observed in yeast (Hillenmeyer et al., 2008).

**Figure 7.**
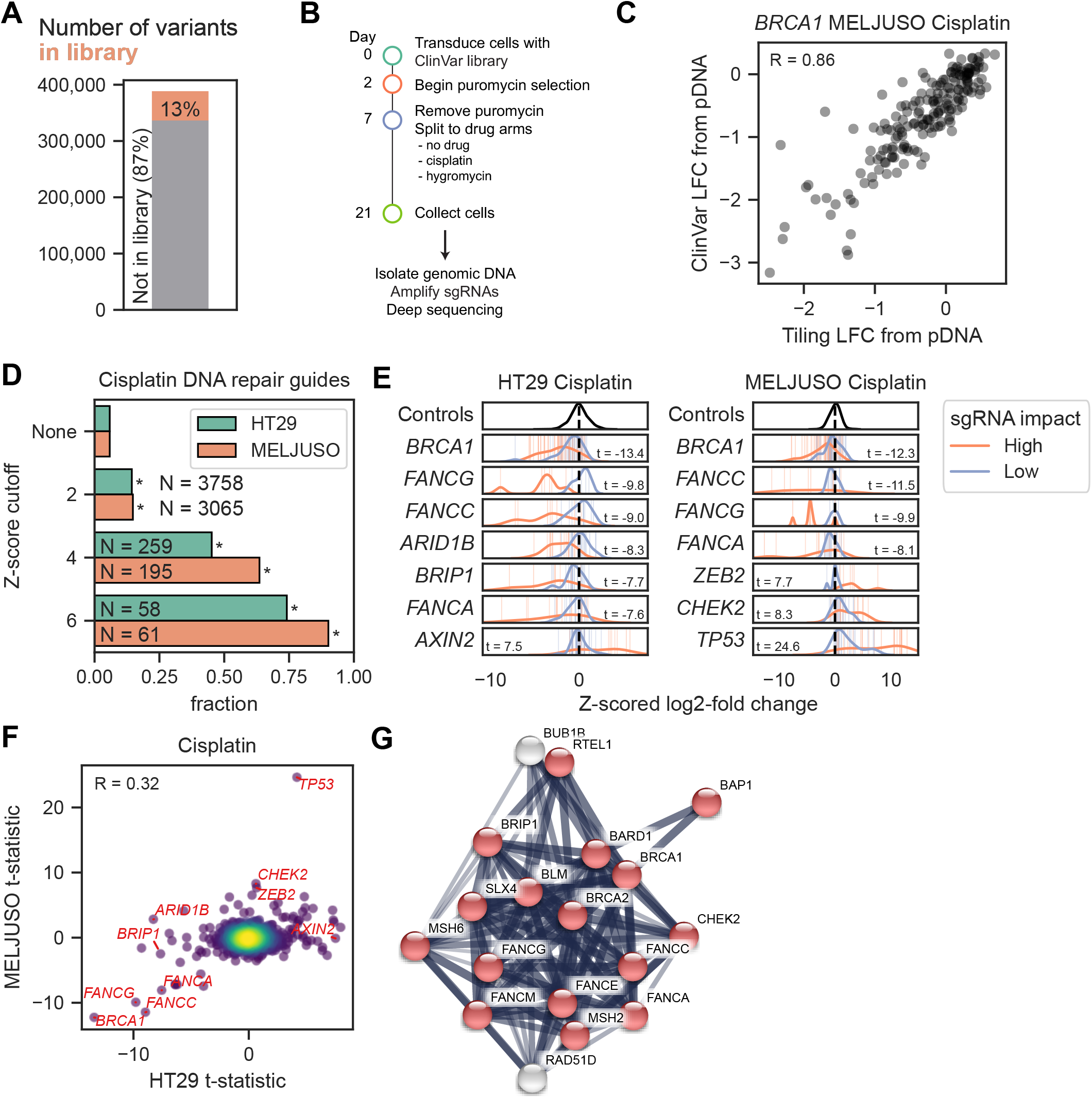
Functional profiling of 52,034 clinical variants. (A) Fraction of all single nucleotide clinical variants (n = 388,496) in the ClinVar library that were included in the library (n = 52,034). (B) Timeline of ClinVar screens in HT29 and MELJUSO cells. (C) Correlation of log-fold changes of sgRNAs targeting BRCA1 between this ClinVar screen and prior tiling screen (Figure 2). Pearson correlation is indicated. (D) Fraction of sgRNAs targeting genes in the Reactome DNA repair pathway at a range of Z-score cutoffs. All cutoffs are significant when compared against the starting library (*p < 1E-16, Fisher’s exact test). (E) Z-scored log-fold change for the top 7 genes with > 3 high and low impact guides in HT29 and MELJUSO cisplatin arms. The mean value for intergenic controls is indicated as a dashed line. Each bar represents a guide, and each density plot represents the distribution of guides targeting a gene. (F) Correlation of the gene-level t-statistic between HT29 and MELJUSO cisplatin arms. The smaller absolute t-statistic when compared against intergenic or non-targeting controls is plotted. Genes from (E) are labeled. (G) Cluster of interactions between hits from HT29 and MELJUSO cisplatin screens. Edges represent confidence in STRING. Nodes colored in red are in the Reactome DNA repair pathway. The complete network can be seen in Figure S8.

Replicates within a cell line were well correlated (Pearson r > 0.84), and we saw good correspondence in log-fold-change values between HT29 and MELJUSO in the untreated arm (Pearson r = 0.9), indicative of many common viability genes, and a lower correlation between cisplatin arms (Pearson r = 0.07), suggesting more cell-line specific differences (**Figure S7B**). Further, log-fold-change values for guides targeting *BRCA1* and *BRCA2* were well correlated between the ClinVar screen and the above-mentioned tiling screens in MELJUSO cells treated with cisplatin (Pearson r = 0.86 for *BRCA1*; Pearson r = 0.78 for *BRCA2*; **Figure 7C; Figure S7C**), demonstrating that large scale base editor screens can yield similar results to more focused libraries. Consistent with the mechanism of cisplatin-induced cytotoxicity, we saw a striking depletion of guides targeting genes in the Reactome-annotated DNA repair pathway in the cisplatin arm (**Figure 7D**). Just 5.8% of sgRNAs in the library targeted DNA repair genes; however, among the sgRNAs that scored with an absolute z-score > 6, that figure increased to 74% for HT29 cells and 90% for MELJUSO cells. Similarly, sgRNAs showing a strong phenotype in the screen were enriched for sgRNAs predicted to introduce pathogenic SNPs. After filtering for guides predicted to introduce a ClinVar-annotated SNP in a non-GC motif and without bystander edits (**Figure S7D**), only 13% (2,626/20,353) of guides were predicted to introduce pathogenic SNPs, but for guides with an absolute z-score > 4, the fraction increased to 42% (28/66) and 44% (25/57) in HT29 and MELJUSO, respectively (**Figure S7E, S7F**). At this cutoff, we identified 24 sgRNAs in HT29 and 11 sgRNAs in MELJUSO that are predicted to introduce variants of unknown significance, representing important guides for future validation.

We next sought to systematically identify genes for which we could effectively assay loss-of-function mutations in these cell line models. To identify significantly enriched genes, we used a two sample t-test comparing the log-fold changes of “high impact” sgRNAs (per VEP annotations; generally nonsense and splice site mutations) to both intergenic and non-targeting controls (**Figure 7E**). Compared with the Kolmogorow-Smirnov (KS) test, the t-test was better at identifying DNA repair genes (KS-test AUC 0.67/0.65 vs t-test AUC 0.75/0.72 for HT29/MELJUSO), suggesting that it is a more sensitive test for these screens, although the two statistical tests largely identified the same genes (Pearson r = 0.87/0.77 for HT29/MELJUSO; **Figure S7G**). Genes behaved similarly across cell lines (Pearson r = 0.32; **Figure 7F**), with *TP53* as the most extreme outlier, showing a much stronger phenotype in MELJUSO, which are *TP53*-wildtype, whereas HT29 cells are *TP53*-mutant.

We used the Benjamini-Hochberg procedure to define gene hits for cisplatin arms using an FDR cutoff of 0.1, obtaining 75 hits total (13 from MELJUSO only, 45 from HT29 only and 17 in both). In agreement with the guide level results, 21 (28%) of these genes are in the Reactome DNA repair pathway, a significant enrichment from the baseline fraction of 39 of 993 (3.9%, including only genes with more than one high impact guide, p-value = 2.2e-9, Fisher’s exact test). Accordingly, we saw a tight cluster of DNA repair genes when we queried the STRING database (Szklarczyk et al., 2019) (**Figure 7G; Figure S8**). We also saw a significant enrichment for genes listed in the GO-term for chromatin organization with 32 of 75 hits (43%) compared with a baseline fraction of 83 of 993 (8.4%, p-value = 3.6e-10, Fisher’s exact test). DNA damage has been shown to induce chromatin reorganization (Mehta et al., 2013), providing a rationale for this enrichment. We also observed gene hits not canonically related to DNA repair but supported by previous results from the literature. For example, the overexpression of two microRNAs targeting *AXIN2*, an inhibitor of the Wnt signaling pathway, has been shown to provide resistance to cisplatin treatment (Chen et al., 2019). In our screen, sgRNAs predicted to introduce high impact variants in *AXIN2* similarly provided resistance to cisplatin treatment in HT29 (t-statistic = 7.5). *LDLR* also scored as a sensitizer to cisplatin treatment (t-statistic = −6.4 in HT29 and −7.2 in MELJUSO); although *LDLR* is best known for its role in cholesterol metabolism, it has recently been shown to sensitize epithelial ovarian cells to cisplatin (Chang et al., 2020). This shows that our screening approach can uncover genes beyond those that are well known to be involved in DNA damage repair.

We saw no correlation for the log-fold changes between HT29 and MELJUSO hygromycin arms (Pearson r = 0.01), reflecting the sparsity of hits in these screens (**Figure S9A**). We did, however, observe *GJB2* as a strong hit in HT29 cells (t-statistic = 9.0), where sgRNAs targeting *GJB2* conferred resistance to hyrgomycin treatment (**Figure S9B**). *GJB2* codes for a structural component of gap junctions, and mutations are implicated in non-syndromic hearing loss (Kelsell et al., 1997). Loss of gap junctions has previously been shown to provide resistance to hygromycin and neomycin, providing mechanistic support for this hit (Yao et al., 2010). Enrichment was not observed in the MELJUSO arm, likely because *GJB2* is not expressed in these cells (log2(TPM + 1) = 0.01 in CCLE).

To determine how the variants introduced by each base editor guide compare to a putative null allele, we conducted a counter-screen with the same library with unmodified Cas9 (WT) in HT29 cells. We binned sgRNAs based on how they performed in the two screens, identifying those that scored in neither screen (BE/WT: |z-score| < 2), both screens (BE/WT: z-score > 4 or < −4), BE only (BE: |z-score| > 4, WT: |z-score| < 2), and WT only (WT: |z-score| > 4, BE: |z-score| < 2) (**Figure S10A-B**). As expected, the BE-only category enriched for VEP high impact edits (**Figure S10C-D**). Some of these guides (34.6%, or 17/49, in dropout; 14.2%, or 11/77, in cisplatin) are positioned such that the base editor modifies a splice acceptor site whereas the cutsite of the wtCas9 nuclease is >10 nts into the intron (**Figure S10E-F**), and thus are a pseudo-false negative of the wtCas9 arm.

For the remaining sgRNAs that score only in the BE arm, an intriguing possibility is the introduction of gain-of-function mutations. To explore this, we identified sgRNAs that (1) scored in only the base editor arm, (2) were categorized as “moderate impact” by the VEP, and (3) did not span an intron-exon boundary; these filters identified 6 sgRNAs in the dropout arm and 21 sgRNAs in the cisplatin arm. The sgRNA with the largest difference between BE and WT screens in the dropout arm targeted *TUBB4A*, which encodes for tubulin β-4A (Z-score: −0.5 WT, −6.4 BE). The predicted edit of this sgRNA, G244S, alters a protein domain that interacts with the GTP molecule at the N-terminal side of the α-tubulin, disrupting tubulin assembly (Hamilton et al., 2014). This variant is categorized in ClinVar as pathogenic, and heterozygous mutations of G244 lead to clinical manifestations in patients with leukodystrophy (Hamilton et al., 2014), supporting a dominant negative mechanism, although validation will be needed to prove this hypothesis. Overall, conducting a counterscreen with Cas9 nuclease provided insight for interpretation of the BE screens, and will be a useful strategy for future screens.

## DISCUSSION

Determining the functional consequences of human genetic variation represents a major challenge that requires scalable technologies. Here, we demonstrate that cytosine base editors can be used to introduce and functionally assess tens of thousands of genetic variants in a pooled screen. We show that deep coverage of individual genes with tiling libraries can be used to identify gain- and loss-of-function variants, including those that modify response to small molecule inhibitors. Further, we show that variants across many genes can be screened in parallel to determine their contribution to a common phenotype, such as sensitivity or resistance to a DNA damaging agent.

The two scales of screens we conduct here can be employed sequentially, whereby a many-gene screen (with a lower density of edits per genes) is first used to identify what genes can be productively examined with base editor technology, allowing subsequent tiling screens of those genes. Likewise, such an approach could be used to nominate genes or domains for saturation mutagenesis experiments, which allow full saturation of all possible single nucleotide variants but come at a substantial cost, rendering them impractical for very large genes or many-gene experiments. Together, these approaches should facilitate the creation of look-up tables that connect gene sequence variations and gene functions, even if a variant has yet to be observed clinically. Furthermore, libraries focused on specific classes of genes may be useful in the drug discovery process. For example, a base editor library that introduces specific mutations into catalytic domains of potential drug targets may be a better surrogate for small molecule inhibition than either knockout or knockdown approaches. Likewise, for small molecules that arise from phenotypic screens, candidate-focused base editor libraries for individual genes or gene families can greatly accelerate the identification of a resistance allele, which remains the gold standard for target identification.

In contrast to conventional screens that focus on the gene as the unit of information, CRISPR-based variant screens present unique challenges. CRISPR knockout, activation, and interference libraries typically include multiple optimized sgRNAs per gene, which mitigates both false positives and false negatives. In base editor screens, however, a given edit can typically be created by a single or small number of sgRNAs; even in cases where multiple sgRNAs are predicted to make the same edit, they are not truly “independent,” as they necessarily have substantial sequence overlap and therefore may have a similar off-target profile. These considerations place additional importance on the validation step. Indeed, as we show, phenotypes may arise from less-expected edits, especially in the context of positive selection screens. Further, because multiple edits can be created by one sgRNA, many genotypes may underlie the ultimate phenotype; thus, the validation process must also include a determination of which edit is causal. Particular care must be taken to avoid over-interpretation of primary base editor screening results, especially for genes with known implications for human health (Gelman et al., 2019).

A key advantage of base editor screens is that they are highly modular, and therefore stand to benefit from innovations in both base editing technology and pooled screening approaches. Here we used CBE technology with canonical SpCas9, but the potential to mix and match alternative base editing domains and Cas proteins promises to expand the space of variants that can be generated with this screening approach. Initial screening attempts using adenine base editors (ABEs), based on the ABE7 architecture (Gaudelli et al., 2017; Koblan et al., 2018), have not yet been productive in our hands, which is consistent with a recent head-to-head comparison of CBEs and ABE7.10 (Kluesner et al., 2020); however, the recent development of high-activity adenine base editors, such as ABE8e and other ABE8 variants (Gaudelli et al., 2020; Richter et al., 2020), may allow the screening workflow developed here to be used with ABEs, increasing the types of nucleotide changes that can be introduced. In addition, base editors that use natural or engineered Cas variants with alternative PAM preferences have been reported (Kim et al., 2017; Kleinstiver et al., 2019; Miller et al., 2020; Nishimasu et al., 2018; Walton et al., 2020), increasing the density of editable target sites, although how these variants perform in a screening setting remains to be determined.

Especially as the depth and types of mutations increases, another potential future direction is to merge base editing screens with combinatorial technologies (Han et al., 2017; Horlbeck et al., 2018; Najm et al., 2017; Sanson et al., 2019) to enable screens with multiple edits per cell. In the same way that synthetic lethal and buffering relationships can be identified with combinatorial loss-of-function screens, combinatorial base editor screens of variants implicated via genome-wide association studies may help to organize variants underlying common diseases with complex genetic underpinnings, although in many cases the availability of cell based models represents an additional experimental hurdle. Likewise, because base editor screens require only a short sequencing read, they are theoretically compatible with alternative pooled screening readouts, such as single-cell RNA sequencing (Datlinger et al., 2017; Dixit et al., 2016; Hill et al., 2018; Replogle et al., 2020) and pooled optical screens (Feldman et al., 2019). In sum, the results presented here establish that base editor screens are a flexible, scalable method to functionally profile variants, and we anticipate that a similar framework will extend pooled variant screening to include a broad array of base editing reagents in a variety of models of human disease.

## ACKNOWLEDGMENTS

We thank Amy Goodale, Edith Sawyer, Jonah O’Mara Schwartz, Hinako Kawabe, Briana Fritchman, and Xiaoping Yang for producing guide libraries and lentivirus; Olivia Bare, Max Macaluso, and Yenarae Lee for operations support; Matthew Greene, Adam Brown, Doug Alan, Mark Tomko, and Tom Green for software engineering support; David Root for senior leadership of the Genetic Perturbation Platform; the Broad Institute Genomics Platform Walk-up Sequencing group for Illumina sequencing.

We thank Greg Findlay (Shendure Lab, University of Washington), Sarah Weiss (Sharpe Lab, Harvard Medical School), Nicky Persky (Genetic Perturbation Platform, Broad Institute), and Kendell Clement and Luca Pinello (Pinello Lab, Massachusetts General Hospital) for helpful conversations.

Schematics were created with BioRender.com. Funding support was provided in part by the Broad V2F Initiative and the Functional Genomics Consortium. LWK and DRL recognize support from the NSF GRFP fellowship (DGE1144152) and NIH U01AI142756.

## AUTHOR CONTRIBUTIONS

Conceptualization: REH, JGD

Investigation: REH, CRF, PCD, AKS, ZMS, ALG, MNF, KRS, YB

Data Curation: REH, MH

Formal Analysis: REH, MH, PCD

Visualization: REH, MH, PCD, ZMS

Resources: LWK, DRL

Supervision: JTN, JGD

Writing - Original Draft: REH, PCD, JGD

Writing - Review & Editing: CRF, LWK, JTN, DRL

Funding Acquisition: JTN, JGD

## METHODS

### Vectors

pRDA_077 (BE3): U6 promoter expresses customizable guide RNA; core EF1a (EFS) expresses codon-optimized BE3 with 2xSV40NLS (Komor et al., 2016) and 2A site provides puromycin resistance. Later sequencing of this vector revealed an E1150K mutation in SpCas9; we therefore constructed pRDA_256 as our preferred BE3 vector for further screens. The pRDA_077 vector was used for screens conducted with the preliminary tiling library (**Figure 1, S1**), the drug resistance tiling library (**Figure 6**), and the ClinVar library (**Figures 7, S7, S8, S9, S10**); for all other screens we used pRDA_256, which restored the wtCas9 sequence. Direct comparison of pRDA_077 and pRDA_256 showed comparable editing efficiency.

pRDA_256 (BE3, Addgene *will be deposited post-COVID*): U6 promoter expresses customizable guide RNA with a 10x guide capture sequence at the 3’ end of the tracrRNA to facilitate future use with direct capture single cell RNA sequencing (Replogle et al., 2020); core EF1a (EFS) expresses codon-optimized BE3 with 2xSV40NLS and 2A site provides puromycin resistance.

pRDA_078 (BE4): U6 promoter expresses customizable guide RNA; core EF1a (EFS) expresses codon-optimized BE4 with 2xSV40NLS (“BE4max”) (Koblan et al., 2018), 2A site provides puromycin resistance.

LentiCRISPRv2 (pXPR_023; Addgene 52961): U6 promoter expresses customizable guide RNA; core EF1a (EFS) promoter expresses wild-type SpCas9 and 2A site provides puromycin resistance.

pLX_311-Cas9 (Addgene 96924): SV40 promoter expresses blasticidin resistance; EF1a promoter expresses wild-type SpCas9.

pRDA_085 (Addgene *will be deposited post-COVID*): EF1a expresses wtCas9; T2A site provides blasticidin resistance and P2A site provides mKate2.

pRDA_118 (modified lentiGuide, Addgene 133459): U6 promoter expresses customizable sgRNA; EF1a promoter provides puromycin resistance. This vector is a derivative of the lentiGuide vector, with a modification to the tracrRNA to eliminate a run of four thymidines.

pMT025 (Addgene *will be deposited post-COVID*): SV40 provides puromycin resistance; EF1a expresses a custom open reading frame.

pLX_313-EGFP (Addgene *will be deposited post-COVID*): SV40 provides hygromycin resistance; EF1a expresses EGFP.

### Tiling library design and annotation

Guide sequences for tiling libraries were designed using sequence annotations from Ensembl (Cunningham et al., 2019). We used Ensembl’s REST API (https://rest.ensembl.org/) to obtain the genomic locations of transcripts, transcript sequences, and protein sequences and used these to annotate each sgRNA with its predicted edits. We included all sgRNAs targeting coding sequence; for all tiling libraries except the preliminary tiling library, we also included all sgRNAs for which the start was up to 30 nucleotides into the intron and UTRs. For the drug resistance tiling library (screened with PARPi), we also identified 14 substitution mutations in 4 genes (*BRAF, EGFR, MAP2K1*, and *PIK3CA*) that appeared in TCGA with >1% frequency (accessed 2019-02-15) and added all possible sgRNAs targeting the mutant alleles (n = 25 sgRNAs). The

For the preliminary tiling library, we included all possible sgRNAs tiling 10 pan-lethal genes (*EEF2, HNRNPU, KPNB1, PELP1, POLR1C, PSMA6, RPS20, SF3B1, SNRPD1*, and *TFRC*), 4 cell surface markers (*CD33, CD81, FAS*, and *ICAM1*), 4 vemurafenib-resistance genes (*CUL3, MED12, NF1*, and *NF2*), and 33 other genes not used in the screens described here; we also included 1,000 non-targeting controls (no targets in the genome). For the drug resistance tiling library (screened with PARPi), we included all possible sgRNAs tiling 19 genes, including *PARP1*, 500 non-targeting sgRNAs, and 482 sgRNAs that target only a single non-gene site (“intergenic”). We additionally designed all possible sgRNAs targeting splice donor sites in essential and nonessential genes (Hart et al., 2014, 2015) and randomly selected 500 of each. The focused tiling libraries for *BRCA1, BRCA2, MCL1*, and *BCL2L1* contained all possible sgRNAs targeting the gene of interest as well as 75 non-targeting controls, 75 intergenic controls, and 32 positive controls targeting splice donor sites in pan-lethal genes.

For the analysis of *BRCA1* and *BRCA2* screens, we also annotated sgRNAs that were predicted to produce edits that were listed in ClinVar. ClinVar annotations (variant_summary.txt) were accessed from the website on October 1,2019 and were filtered to include only variants with a review status of at least one gold star. We only annotated sgRNAs if the ClinVar SNP was an exact match to the mutation predicted by the sgRNA (i.e. same nucleotide change and, for coding regions, same amino acid change).

To obtain a “mutation bin” for each sgRNA, we ordered the mutation types as: Nonsense > Splice site > Missense > Intron > Silent > UTR. Guides containing multiple mutation types were binned as the most severe mutation type. Guides predicted to make no edits in the editing window were binned as “No edits.” Likewise, to obtain a “clinical significance bin,” we classified sgRNAs predicted to introduce multiple ClinVar SNPs based on the most severe clinical significance: Pathogenic > Likely pathogenic; Pathogenic / Likely pathogenic > Uncertain significance > Conflicting reports of pathogenicity > Variant not listed in ClinVar > Likely benign; Benign / Likely benign > Benign. With this ordering, sgRNAs were only binned as “Likely benign” or “Benign” if they did not introduce any mutations not listed in ClinVar, which effectively have an unknown functional significance.

### ClinVar library design

The ClinVar library was designed to contain all possible sgRNAs targeting SNPs found in the ClinVar database (accessed 2019-03-14) that are editable with the C>T base editor in the strict window of position 4-8 along the sgRNA. Here, we included sgRNAs predicted to make edits in GC motifs. The same 1982 control sgRNAs as described above in the drug resistance library design were included. The library was filtered to exclude any sgRNAs with BsmBI sites or a TTTT sequence resulting in a final library of 68,526 sgRNAs spanning 52,034 SNPs and 3,584 genes. The library was cloned into pRDA_077 (BE3).

### Annotation of ClinVar library using Ensembl Variant Effect Predictor

The REST API (https://rest.ensembl.org/#VEP, v12.0) of the Ensembl Variant Effect Predictor (VEP) (McLaren et al., 2016) was used to annotate the bystander edits of sgRNAs in the ClinVar library. For every sgRNA, the input to VEP was the chromosome, genomic position of start and end of the edit window, and the edited window sequence; for single edits, only the position of the edit itself was included. We also extracted the APPRIS, Transcript Support Level (TSL) and MANE annotations for each transcript. To pick one relevant transcript annotation for each guide, we first obtained all the transcripts whose “consequence terms” matched the “most severe consequence” for that input. The remaining transcripts were then ranked based on their “Source annotation” (HGNC > EntrezGene > Clone_based_ensembl_gene), APPRIS annotation and TSL.

### Analysis of ClinVar screens

To condense ClinVar annotations, we mapped each clinical variant to its expected phenotype (i.e. likely benign to benign, pathogenic/likely pathogenic to pathogenic etc.). Then for analyzing clinical variant annotations, we only considered sgRNAs creating a single edit in a non-GC motif.

For VEP consequences, we mapped sgRNAs to their most severe predicted consequence as ranked by Ensembl. We then annotated sgRNAs which have all C’s in a GC-motif in the edit window as “No edit”. Control sgRNAs were annotated separately as “negative controls” - sgRNAs targeting non-essential splice sites, intergenic and non-targeting controls - and “positive controls” - sgRNAs targeting essential splice sites.

To determine the significance of genes, we used a t-test, comparing the distribution of VEP high impact guides targeting each gene to either set of negative controls: non-targeting and intergenic guides. The smaller (less-siignificant) absolute t-statistic is reported.

### Library production

Oligonucleotide pools were synthesized by CustomArray. BsmBI recognition sites were appended to each sgRNA sequence along with the appropriate overhang sequences (bold italic) for cloning into the sgRNA expression plasmids, as well as primer sites to allow differential amplification of subsets from the same synthesis pool. The final oligonucleotide sequence was thus: 5’-[Forward Primer]CGTCTCA***CACCG***[sgRNA, 20 nt]***GTTT***CGAGACG[Reverse Primer]. Primers were used to amplify individual subpools using 25 μL 2x NEBnext PCR master mix (New England Biolabs), 2 μL of oligonucleotide pool (~40 ng), 5 μL of primer mix at a final concentration of 0.5 μM, and 18 μL water. PCR cycling conditions: 30s at 98°C, 30s at 53°C, 30s at 72°C, for 24 cycles. In cases where a library was divided into subsets unique primers could be used for amplification:

Primer Set; Forward Primer, 5’ - 3’; Reverse Primer, 5’ - 3’
1; AGGCACTTGCTCGTACGACG; ATGTGGGCCCGGCACCTTAA
2; GTGTAACCCGTAGGGCACCT; GTCGAGAGCAGTCCTTCGAC
3; CAGCGCCAATGGGCTTTCGA; AGCCGCTTAAGAGCCTGTCG
4; CTACAGGTACCGGTCCTGAG; GTACCTAGCGTGACGATCCG
5; CATGTTGCCCTGAGGCACAG; CCGTTAGGTCCCGAAAGGCT
6; GGTCGTCGCATCACAATGCG; TCTCGAGCGCCAATGTGACG

The resulting amplicons were PCR-purified (Qiagen) and cloned into the library vector via Golden Gate cloning with Esp3l (Fisher Scientific) and T7 ligase (Epizyme); the library vector was pre-digested with BsmBI (New England Biolabs). The ligation product was isopropanol precipitated and electroporated into Stbl4 electrocompetent cells (Life Technologies) and grown at 30 °C for 16 h on agar with 100 μg mL-1 carbenicillin. Colonies were scraped and plasmid DNA (pDNA) was prepared (HiSpeed Plasmid Maxi, Qiagen). To confirm library representation and distribution, the pDNA was sequenced.

### Lentivirus production

For small-scale virus production, the following procedure was used: 24 h before transfection, HEK293T cells were seeded in 6-well dishes at a density of 1.5 × 10^6^ cells per well in 2 mL of DMEM + 10% FBS. Transfection was performed using TransIT-LT1 (Mirus) transfection reagent according to the manufacturer’s protocol. Briefly, one solution of Opti-MEM (Corning, 66.25 μL) and LT1 (8.75 μL) was combined with a DNA mixture of the packaging plasmid pCMV_VSVG (Addgene 8454, 250 ng), psPAX2 (Addgene 12260, 1250 ng), and the transfer vector (e.g., pLentiGuide, 1250 ng). The solutions were incubated at room temperature for 20-30 min, during which time media was changed on the HEK293T cells. After this incubation, the transfection mixture was added dropwise to the surface of the HEK293T cells, and the plates were centrifuged at 1000 g for 30 min at room temperature. Following centrifugation, plates were transferred to a 37 °C incubator for 6-8 h, after which the media was removed and replaced with DMEM +10% FBS media supplemented with 1% BSA.

A larger-scale procedure was used for pooled library production. 24 h before transfection, 18 × 10^6^ HEK293T cells were seeded in a 175 cm^2^ tissue culture flask and the transfection was performed the same as for small-scale production using 6 mL of Opti-MEM, 305 μL of LT1, and a DNA mixture of pCMV_VSVG (5 μg), psPAX2 (50 μg), and 40 μg of the transfer vector. Flasks were transferred to a 37 °C incubator for 6-8 h; after this, the media was aspirated and replaced with BSA-supplemented media. Virus was harvested 36 h after this media change.

### Cell culture

A375, MELJUSO, OVCAR8, and HT29 cells were obtained from Cancer Cell Line Encyclopedia and HA1E cells were obtained from the Connectivity Map, both at the Broad Institute. HAP1 cells were obtained from Horizon Discovery (item C631). HEK293Ts were obtained from ATCC (CRL-3216). All cells regularly tested negative for mycoplasma contamination and were maintained in the absence of antibiotics except during screens, validation experiments, and lentivirus production, during which media was supplemented with 1% penicillin-streptomycin. Cells were passaged regularly (every 2-4 days) to maintain exponential growth and were kept in a humidity-controlled 37°C incubator with 5.0% CO_2_. Media conditions, and doses of polybrene, puromycin, blasticidin, and hygromycin were as follows:

A375: RPMI + 10% fetal bovine serum (FBS); 1 μg/mL; 1 μg/mL; 5 μg/mL; N/A
HA1E: MEM alpha + 10% FBS; 4 μg/mL; 1 μg/mL; N/A; N/A
HAP1: IMDM + 10% FBS; 4 μg/mL; 1 μg/mL; 5 μg/mL; N/A
HEK293T: DMEM + 10% heat-inactivated FBS; N/A; N/A; N/A; N/A
HT29: DMEM + 10% FBS; 1 μg/mL; 2 μg/mL; 8 μg/mL; N/A
MELJUSO: RPMI + 10% FBS; 4 μg/mL; 1 μg/mL; 4 μg/mL; 100 μg/mL
OVCAR8: RPMI + 10% FBS; 4 μg/mL; 1 μg/mL; 8 μg/mL; N/A

For screens in A375 cells, selumetinib (Selleckchem, S1008) was screened at a dose of 1.5 μM and vemurafenib (Selleckchem, S1267) was screened at 2 μM. For ClinVar screens, cisplatin (BioVision, 10187) was diluted in 0.9% NaCI and was screened at 8 μM in HT29 and 1 μM in MELJUSO. Hygromycin (Life Technologies, 10687010) was screened at 125 μg/mL in HT29 and 32.25 μg/mL in MELJUSO. For *MCL1* and *BCL2L1* screens and subsequent validation in MELJUSO cells, S63845 (referred to as “MCL1-i”; gift from Guo Wei) and A-1331852 (referred to as “BCL2L1-i”; Active Biochem, A-6046) were both screened at 250 nM (in the single inhibitor-treated screens) and 62.5 nM (in the BCL2L1-i and MCL1-i combination-treated screens). For *PARP1* tiling screens and subsequent validation in HAP1 cells, niraparib (Selleckchem, S2741), olaparib (Cayman Chemical, 10621), rucaparib (Selleckchem, S1098), talazoparib (Selleckchem, S7048), and veliparib (Selleckchem, S1004) were screened at doses of 200 nM, 2 μM, 2 μM, 3 nM, and 14 μM, respectively. For *BRCA1* and *BRCA2* screens in MELJUSO cells, cisplatin was screened at 1 μM and talazoparib was screened at 7.81 nM.

### Determination of antibiotic dose

In order to determine an appropriate antibiotic dose for each cell line, cells were transduced with the pRosetta or pRosetta_v2 lentivirus such that approximately 30% of cells were infected and therefore EGFP+. At least 1 day post-transduction, cells were seeded into 6-well dishes at a range of antibiotic doses (e.g. from 0 μg/mL to 8 μg/mL of puromycin). The rate of antibiotic selection at each dose was then monitored by performing flow cytometry for EGFP+ cells. For each cell line, the antibiotic dose was chosen to be the lowest dose that led to at least 95% EGFP+ cells after antibiotic treatment for 7 days (for puromycin) or 14 days (for blasticidin and hygromycin).

### Determination of lentiviral titer

To determine lentiviral titer for transductions, cell lines were transduced in 12-well plates with a range of virus volumes (e.g. 0, 150, 300, 500, and 800 μL virus) with 1 to 1.5 × 10^6^ cells per well in the presence of polybrene. The plates were centrifuged at 640 x g for 2 h and were then transferred to a 37 °C incubator for 4-6 h. Each well was then trypsinized, and an equal number of cells seeded into each of two wells of a 6-well dish. Two days post-transduction, puromycin was added to one well out of the pair. After 5 days, both wells were counted for viability. A viral dose resulting in 30-50% transduction efficiency, corresponding to an MOI of ~0.35-0.70, was used for subsequent library screening.

### Derivation of stable cell lines

In order to establish wtCas9 expressing cell lines for screens with the preliminary tiling library, MELJUSO cells and A375 cells were transduced with pLX_311-Cas9 and pRDA_085, respectively, and successfully transduced cells were selected with blasticidin for a minimum of 2 weeks.

### Pooled screens

For pooled screens, cells were transduced in 2-3 biological replicates with the lentiviral library. Transductions were performed at a low multiplicity of infection (MOI ~0.5), using enough cells to achieve a representation of at least 500 transduced cells per sgRNA assuming a 30-50% transduction efficiency. Because the titer of all-in-one base editor viruses was low, we plated cells in polybrene-containing media with 1 x 10^6^ - 3 x 10^6^ cells per well in a 12-well plate (depending on the infection efficiency observed in the titration). Plates were centrifuged for 2 hours at 640 x g and transferred to an incubator for 4-6 hours, after which cells were pooled into flasks. Puromycin was added 2 days post-transduction and maintained for 5-7 days to ensure complete removal of non-transduced cells. Upon puromycin removal, cells were split to any drug arms (each at a representation of at least 1,000 cells per sgRNA) and passaged every 2-4 days for an additional 2 weeks to allow sgRNAs to enrich or deplete; cell counts were taken at each passage to monitor growth. For ClinVar screens, drug treatment was pulsed twice (3 days on, 4 days off). The wtCas9 screen was terminated after the second drug pulse (17 days post-transduction) due to COVID-19 related lab shutdowns. At the conclusion of each screen, cells were pelleted by centrifugation, resuspended in PBS, and frozen promptly for genomic DNA isolation.

For the screen of the preliminary tiling library in A375 cells stably expressing wtCas9, a no-spin transduction was performed. A mixture of cells, lentivirus, and polybrene at 0.5 μg/mL was divided into T175 flasks and placed in an incubator overnight. Approximately 16 hours later, the lentivirus-containing media was replaced with fresh media and cells were returned to the incubator. After this point, the screen was performed as described above.

### Genomic DNA isolation and sequencing

Genomic DNA (gDNA) was isolated using either the KingFisher Flex Purification System with the Mag-Bind^®^ Blood & Tissue DNA HDQ Kit (Omega Bio-Tek #M6399-01), or the Machery Nagel NucleoSpin Blood Maxi (2e7-1e8 cells), Midi (5e6-2e7 cells), or Mini (<5e6 cells) kits, per the manufacturer’s instructions. The gDNA concentrations were quantitated by Qubit. For samples where genomic DNA was limiting, gDNA was purified prior to PCR using the Zymo OneStep PCR Inhibitor Removal Kit (Zymo, D6030), per the manufacturer’s instructions.

For PCR amplification, gDNA was divided into 100 μL reactions such that each well had at most 10 μg of gDNA. Plasmid DNA (pDNA) was also included at a maximum of 100 pg per well. PCR amplification for base editor screens and A375 wtCas9 screen with the initial essential gene tiling library was performed with Ex Taq (Takara); all other screens were amplified using Titanium Taq (Takara) and 5% DMSO, which we recommend going forward for improved PCR efficiency. Per 96 well plate, a master mix consisted of 150 μL DNA Polymerase (Ex Taq or Titanium Taq; Takara), 1 mL of 10x buffer, 800 μL of dNTPs (Takara), 50 μL of P5 stagger primer mix (stock at 100 μM concentration), 500 μL of DMSO (if used), and water to bring the final volume to 4 mL. Each well consisted of 50 μL gDNA plus water, 40 μL PCR master mix, and 10 μL of a uniquely barcoded P7 primer (stock at 5 μM concentration). PCR cycling conditions were as follows: an initial 1 min at 95 °C; followed by 30 s at 94 °C, 30 s at 52.5 °C, 30 s at 72 °C, for 28 cycles; and a final 10 min extension at 72 °C. PCR primers were synthesized at Integrated DNA Technologies (IDT). PCR products were purified with Agencourt AMPure XP SPRI beads according to manufacturer’s instructions (Beckman Coulter, A63880). Samples were sequenced on a HiSeq2500 HighOutput (Illumina) with a 5% spike-in of PhiX.

### Screen analysis

Guide sequences were extracted from sequencing reads by running the PoolQ tool with the search prefix “CACCG” (https://portals.broadinstitute.org/gpp/public/software/poolq). Reads were counted by alignment to a reference file of all possible guide RNAs present in the library. The read was then assigned to a condition (e.g. a well on the PCR plate) on the basis of the 8 nt index included in the P7 primer. After deconvolution, read counts were log-normalized by the following formula:

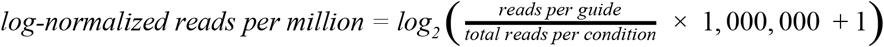

We then calculated the log-fold change between conditions. All dropout (no drug) conditions were compared to the plasmid DNA (pDNA); drug-treated conditions were compared to the time-matched dropout sample, with the exception of MELJUSO cells in the *BRCA1* and *BRCA2* screens, which were compared to the plasmid DNA because loss of *BRCA1* and *BRCA2* had some viability effect in the absence of drug.

Prior to further analysis, we filtered out sgRNAs for which the log-normalized reads per million of the pDNA was > 3 standard deviations from the mean. We also filtered out any sgRNAs containing more than 5 off-target sites in the human genome with a CFD score of 1.0 (indicating a perfect or near-perfect match). The total number of off-targets with a CFD score = 1.0 is provided in the “Match Bin I sum” column in the Supplementary Tables.

### Validation experiments

For validation experiments in which the target site was directly sequenced, individual sgRNAs were cloned into either RDA_077 or RDA_256 and made into lentivirus as described above. At least 3 x 10^6^ cells were transduced in duplicate with a virus volume to obtain ~30-50% transduction efficiency and were selected with puromycin for 5-7 days to remove uninfected cells; puromycin doses were as described above, except for HAP1 cells, which were treated with puromycin at 2 μg/mL. After puromycin selection was removed, cells were split into any drug arms and cultured for an additional 14 days (for *BRCA1, BRCA2, MCL1*, and *BCL2L1* sgRNAs) or 12 days (for *PARP1* sgRNAs). For one replicate of cells containing sg19-21 and co-treated with BCL2L1-i and MCL1-i, cells were cultured for an additional week on drug (3 weeks total) to obtain enough cell material for sequencing.

Genomic DNA was isolated using the Kingfisher as described above, and the target sites were amplified using a 2-step PCR. In the first round of PCR, genomic DNA was amplified using custom primers designed to amplify each target site. Each well contained 50 μL of NEBNext High Fidelity 2X PCR Master Mix (New England Biolabs, M0541), 0.5 μL of each primer at 100 μM, and 49 μL of gDNA. We used a touch-down PCR with the following cycling conditions: (1) 98°C for 1 minute; (2) 98°C for 30 seconds; (3) 68°C for 30 seconds (−1° per cycle); (4) 72°C for 1 minute; (5) Go to step 2, x 15; (6) 72°C for 10 minutes. The second round of PCR appended Illumina adapters and well barcodes for sequencing using the P5 primer “Argon” and the P7 primer “Kermit”. Each well contained 1.5 μL of Titanium Taq (Takara), 10 μL of Titanium Taq buffer, 8 μL of dNTPs, 5 μL of DMSO, 0.5 μL of P5 primer (Argon) at 100 μM, 10 μL of P7 primer (Kermit), 55 μL of water, and 10 μL of PCR product from the first PCR. The following cycling conditions were used: (1) 95°C for 1 minute; (2) 94°C for 30 seconds; (3) 52.5°C for 30 seconds; (4) 72°C for 30 seconds; (5) go to (2), x 15; (6) 72°C for 10 minutes. Each well was separately purified with Agencourt AMPure XP SPRI beads according to the manufacturer’s instructions (Beckman Coulter, A6388O), using a 1:1 ratio of beads to PCR product. DNA concentration was quantified using a Nanodrop and wells were pooled proportionally to their concentrations. The pooled library was quantified by Qubit and sequenced using the Illumina MiSeq with a 300 nucleotide single read and a 10% PhiX spike-in.

### Analysis of deep sequencing data

CRISPResso2 (version 2.0.30) was used to process all sequencing reads from validation experiments (Clement et al., 2019). CRISPResso2 was run in base editor mode using the default settings with the following changes: --exclude_bp_from_left 8 --exclude_bp_from_right 8 --min_average_read_quality 25. We also set custom values for each sgRNA for --plot_window_size, -wc, and --default_min_aln_score. All samples had >1,000 aligned reads and 400/418 samples had >10,000 aligned reads.

To calculate replicate correlations, we used the “Alleles_frequency_table_around_sgRNA” file from the CRISPResso2 output, which contains the read counts for each allele (defined as a subsequence around the sgRNA, the length of which is specified by --plot_window_size). We then log-normalized the read counts for each sample (using the same formula described in the “Analysis of screens” section). Finally, we filtered out any alleles with <100 reads in all replicates and drug conditions for that sgRNA, and calculated the Pearson correlation between log-normalized reads.

For further analysis of alleles, we set a more stringent filter in order to avoid spurious log-fold change values due to low read counts. We filtered out any alleles that comprised < 1% of the total reads in all replicates and drug conditions for that sgRNA.

### Overexpression of mutant cDNAs

The wild-type and S145F mutant versions of the *BCL2L1* CDS were cloned into pMT025 and packaged into lentivirus as described above. Unmodified parental MELJUSO cells were transduced with a virus volume to obtain approximately 30-50% transduction efficiency and transduced cells were selected with puromycin for 8 days. As a control, we also included unmodified parental cells and MELJUSO cells stably expressing pLX_313 (EGFP); the latter cells were selected with hygromycin and confirmed to express EGFP by flow cytometry (BD Accuri). All 4 cell lines were seeded in 96-well plates in quadruplicate and treated with a serial dilution of BCL2L1-i (diluted four-fold from 32 μM to 0.03 nM, plus untreated) and MCL1-i at 3 doses (untreated, 62.5 nM, or 250 nM). Cell viability was assayed 4 days post-seeding by Cell Titer Glo (Promega) and an Envision plate reader, according to the manufacturer’s instructions. Cell Titer Glo values were normalized to cells not treated with BCL2L1-i to determine relative viability for each cell line at each dose of MCL1-i.

### External datasets

Saturation genome editing data for **Figure 2** was accessed from the original publication (Findlay et al., 2018). Genomic coordinates supplied with the dataset were converted to the GRCh38 build using liftOver. The data for the pan-lethal tiling library screened in MELJUSO cells expressing wtCas9 are from (Sanson et al., 2019); raw data are also included here.

### Data visualization

Figures were created with Python and GraphPad Prism; schematics were created with BioRender.com. PyMOL (version 2.3.2) was used to map the screening data onto the following crystal structures from the Protein Data Bank: 5LOF (MCL1 in complex with S63845), 4QVX (BCL2L1 in complex with A-155463), and 4OQB (PARP1 in complex with a DNA double-strand break and PARP inhibitor).

---

**Figure S1.**
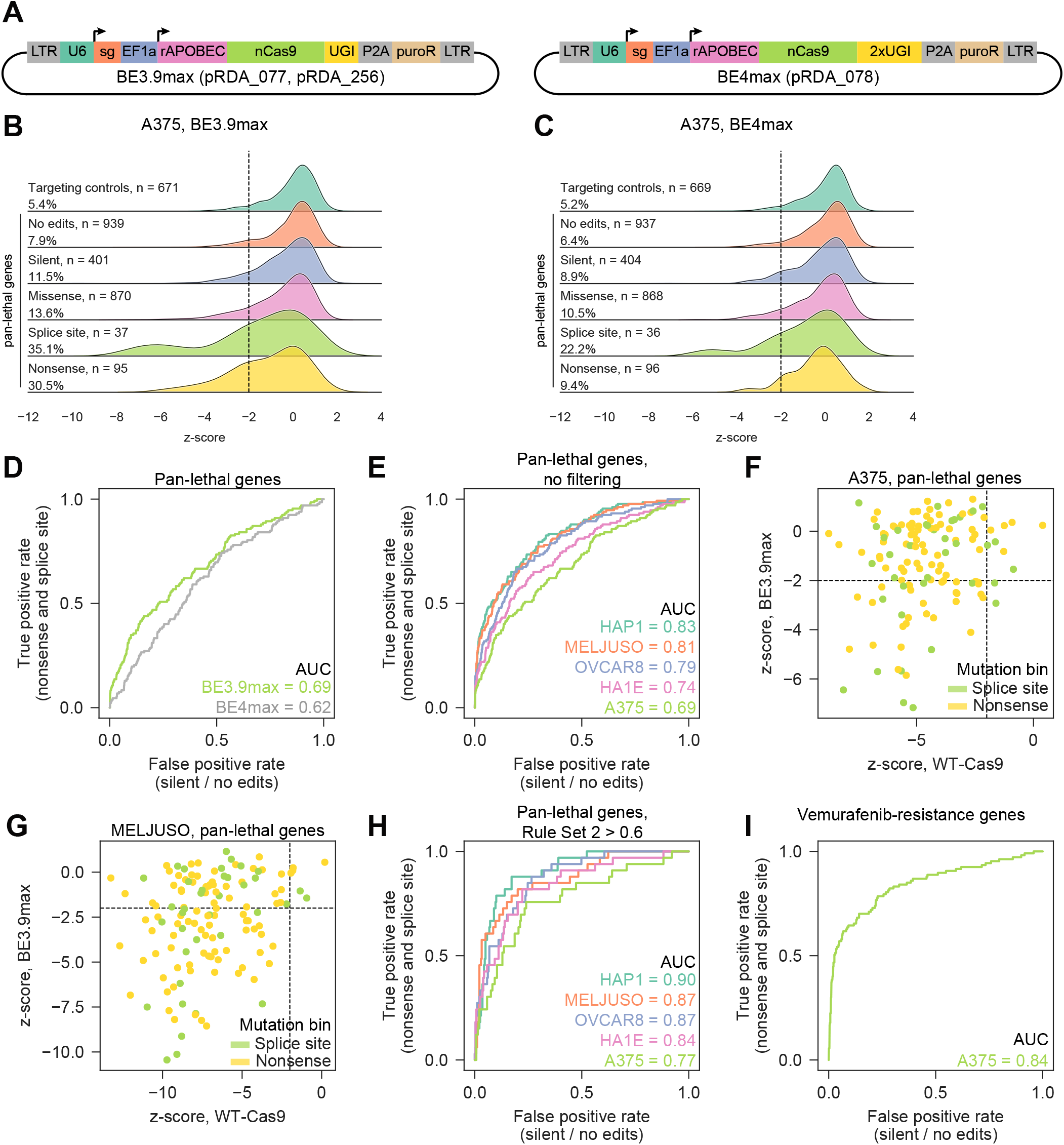
Comparison of BE3.9max, BE4max, and wtCas9 in tiling screens, related to Figure 1. (A) Lentiviral vectors used for base editor screens. (B-C) Performance of sgRNAs targeting pan-lethal genes in A375 screens using BE3.9max (B) and BE4max (C). Guides are grouped by the most severe predicted mutation consequence; the percentage of sgRNAs in each mutation bin with a z-score < −2 is also shown. Targeting controls are all sgRNAs targeting cell surface markers, regardless of mutation consequence. The set size differs slightly between BE3.9max and BE4max due to removal of sgRNAs with outlier abundance in the plasmid DNA. (D) Receiver operating characteristic (ROC) curves for sgRNAs targeting pan-lethal genes in screens with BE3.9max and BE4max. True positives: sgRNAs predicted to introduce nonsense and splice site mutations (n = 132). True negatives: sgRNAs predicted to introduce no edits or silent mutations (n = 1340 for BE3.9max; n = 1341 for BE4max). (E) ROC curves for sgRNAs targeting pan-lethal genes in screens with BE3.9max in 5 cell lines. True positives and true negatives as in (D); n = 132 true positives; n = 1340 true negatives. (F-G) Performance of sgRNAs introducing nonsense or splice site mutations in pan-lethal genes in screens with wtCas9 or BE3.9max, for A375 (F) and MELJUSO (G). Pearson r = 0.22 for A375; Pearson r = 0.28 for MELJUSO; n = 132 sgRNAs. Dotted line on each axis indicates a z-score of −2. (H) ROC curves for sgRNAs targeting pan-lethal genes in screens with BE3.9max in 5 cell lines; only sgRNAs with a Rule Set 2 score > 0.6 are included. True positives and true negatives as in (D) and (E); n = 33 true positives; n = 279 true negatives. (I) ROC curve for sgRNAs targeting *NF1, NF2, MED12*, and *CUL3* in vemurafenib-resistance screens in A375. True positives and true negatives as in (D), (E), and (H); n = 107 true positives; n = 945 true negatives.

**Figure S2.**
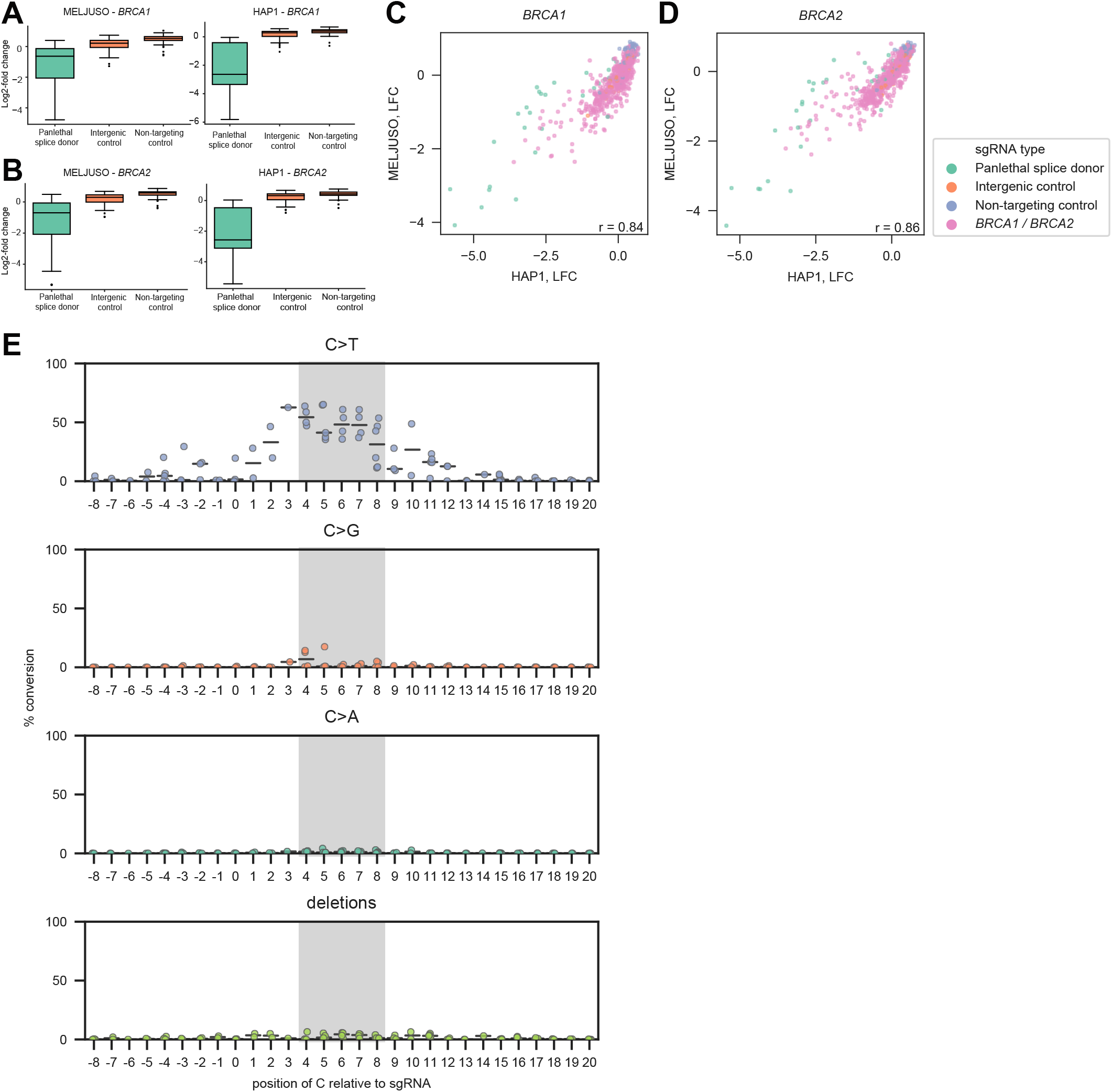
Base editor screens of *BRCA1* and *BRCA2* identify known loss-of-function mutations, related to Figure 2. (A) Distribution of negative control sgRNAs (n = 74 non-targeting sgRNAs; n = 74 intergenic sgRNAs) and positive control sgRNAs (n = 31 sgRNAs targeting splice sites in pan-lethal genes) for BRCA1 screens in HAP1 and MELJUSO cells (in the no drug arms). Boxes show the quartiles; whiskers show 1.5 times the interquartile range. (B) Distribution of negative control sgRNAs (n = 75 non-targeting sgRNAs; n = 74 intergenic sgRNAs) and positive control sgRNAs (n = 31 sgRNAs targeting splice sites in pan-lethal genes) for BRCA2 screens in HAP1 and MELJUSO cells (in the no drug arms). Boxes show the quartiles; whiskers show 1.5 times the interquartile range. (C-D) Correlation between the log-fold change (LFC) of HAP1 cells (no drug arm) and MELJUSO cells (average of talazoparib- and cisplatin-treated arms) for *BRCA1* screen (C) and *BRCA2* screen (D). Pearson r is reported. Colors indicate the sgRNA target. (E) Percentage of conversion at cytosines based on the position within the protospacer, for the 13 sgRNAs validated for *BRCA1* and *BRCA2*. The position of the C is indicated on the x-axis, where 1 corresponds to the first nucleotide of the protospacer and 21-23 correspond to the PAM. The expected window of editing (4-8) is shown in gray.

**Figure S3.**
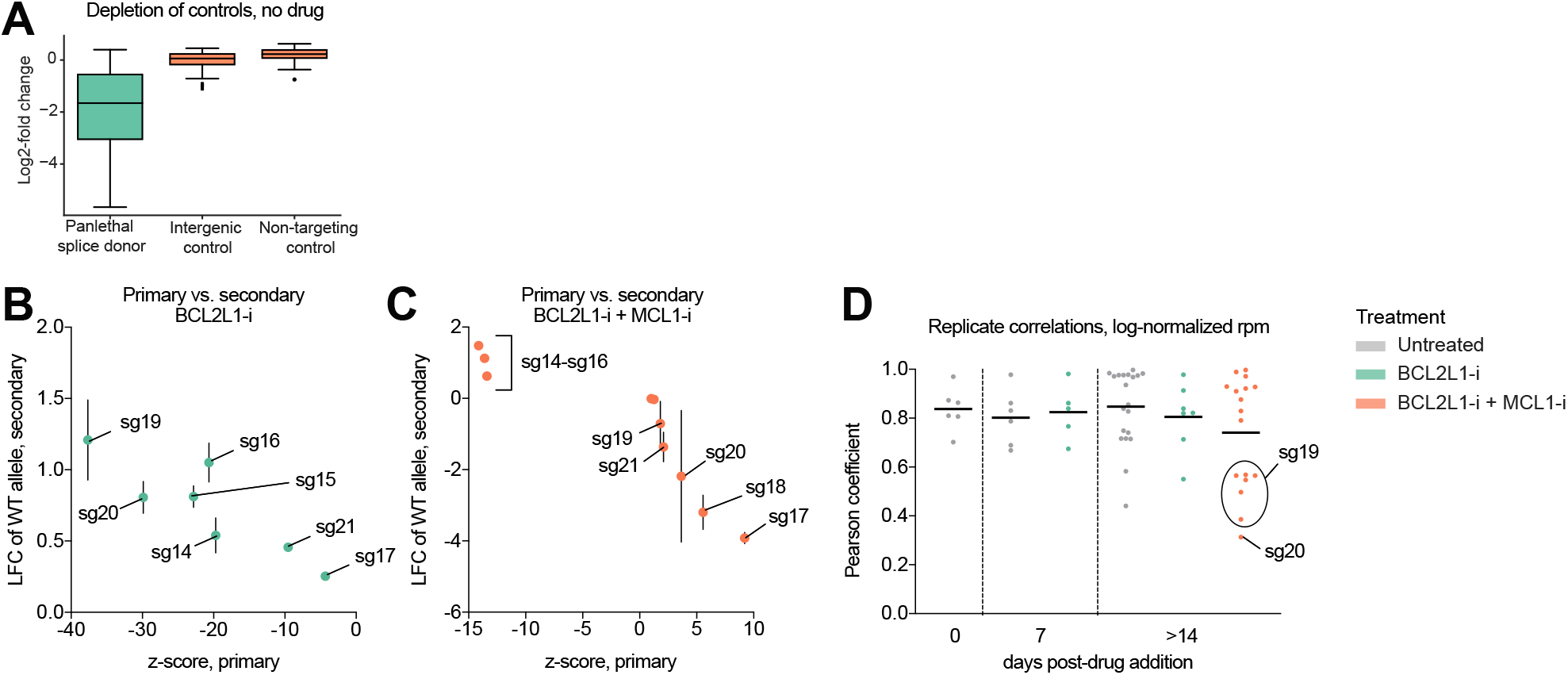
Quality control metrics for *MCL1* screen and validation, related to Figure 4. (A) Distribution of negative control sgRNAs (n = 75 non-targeting sgRNAs; n = 75 intergenic sgRNAs) and positive control sgRNAs (n = 32 sgRNAs targeting splice sites in pan-lethal genes) for *MCL1* tiling screen. Boxes show the quartiles; whiskers show 1.5 times the interquartile range. (B) Comparison between sgRNA performance in the primary screen (z-score) and secondary validation (LFC of the wild-type allele at 14 days post-drug addition), for BCL2L1-i treated cells. Error bars show the range of n = 2 biological replicates in the secondary validation. (C) Comparison between primary screen and secondary validation as in (B), for BCL2L1-i and MCL1-i co-treated cells. Error bars show the range of biological replicates in the secondary validation. N = 1 replicate for sg14-sg16; n = 2 replicates for sg17, sg18, sg20, and sg22-sg23; n = 4 replicates for sg19 and sg21. (D) Distribution of Pearson coefficients for allele-level replicate correlations for sg14-23 at each drug condition and timepoint. The mean is shown in black.

**Figure S4.**
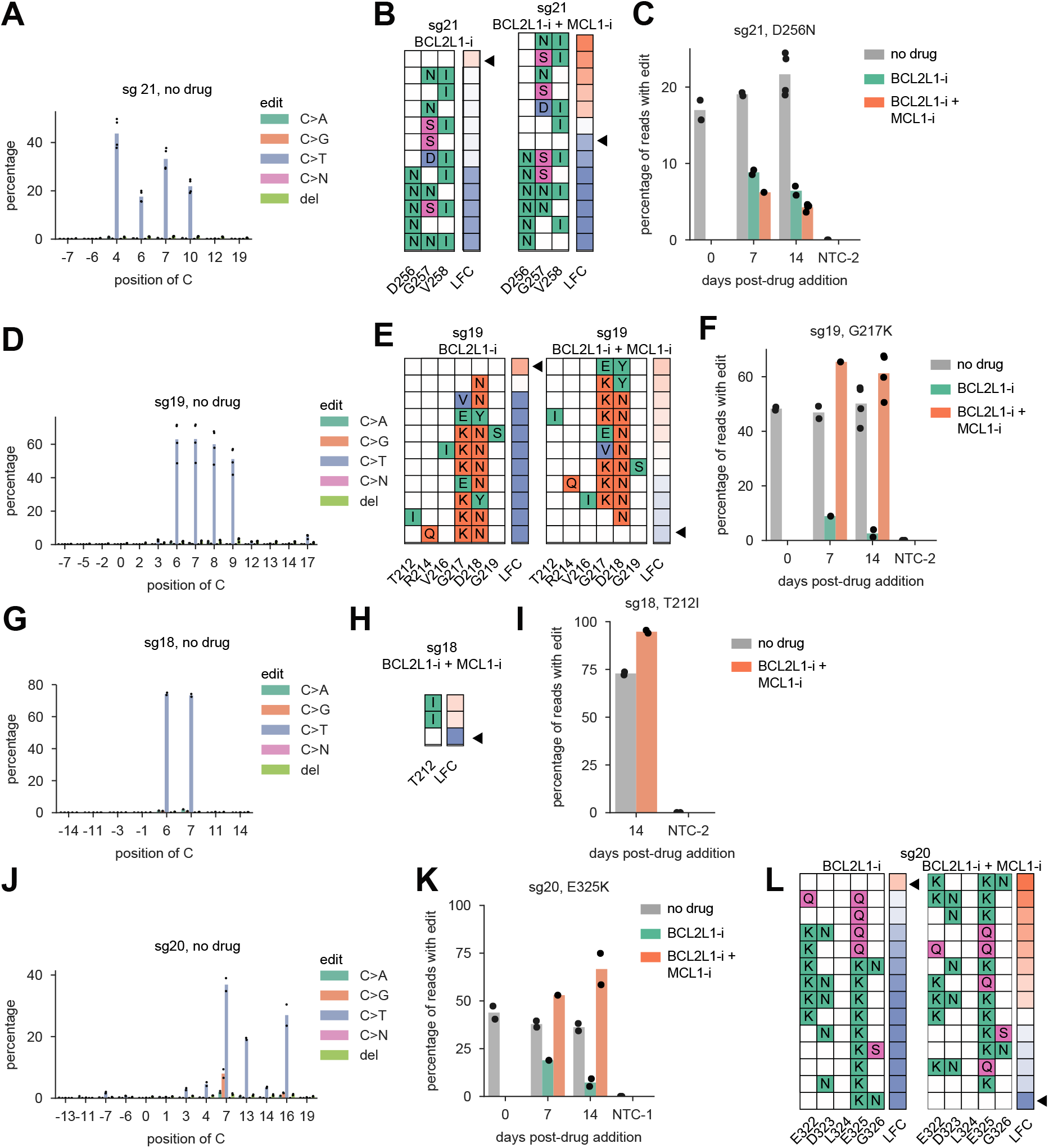
Validation of mutations in MCL1 that confer resistance and sensitivity to MCL1-i, related to Figure 4. (A) In untreated cells, sg21 shows efficient C>T editing at C4, C6, C7 and C10. Cytosines are numbered by position in the sgRNA, where 1 is the first nucleotide of the protospacer and 21-23 is the PAM. Dots show n = 4 biological replicates. (B) Alleles containing a D256N mutation are depleted in both BCL2L1-i-treated cells and BCL2L1-i, MCL1-i co-treated cells relative to untreated cells. Conversely, alleles containing G257N and G257S mutations enriched upon BCL2L1-i and MCL1-i co-treatment. The triangle indicates the wild-type (unedited) allele. (C) Percentage of all sequencing reads containing the indicated edit for each timepoint and drug condition. Reads containing D256N mutations depleted from 21.6% in untreated cells to 4.3% in BCL2L1-i and MCL1-i co-treated cells. Dots show n = 4 biological replicates. Reads containing indels were not considered to be edited. NTC-2 indicates an untreated, non-targeting control (EGFP-targeting sgRNA), sequenced at 14 days post-drug addition. (D) Pattern of base editing for sg19 in untreated cells. Dots show n = 4 biological replicates. (E) All edited alleles deplete in BCL2L1-i-treated cells relative to untreated cells, whereas edited alleles enrich slightly in BCL2L1-i and MCL1-i co-treated cells. (F) The G217K mutation depleted strongly from 50.2% of reads in untreated cells to 2.6% of reads in BCL2L1-i-treated cells, but enriched slightly to 61.3% of reads in co-treated cells. (G) Pattern of base editing for sg18 in untreated cells. Dots show n = 2 biological replicates. (H) Alleles containing a T212I mutation are enriched in cells co-treated with BCL2L1-i and MCL1-i relative to untreated cells. (I) Reads with T212I edit enrich from an average of 72.9% in untreated cells to an average of 94.8% in BCL2L1-i and MCL1-i co-treated cells. (J) Pattern of base editing by sg20. Dots show n = 2 biological replicates. (K) Reads containing the E325K mutation depleted from 36.3% in untreated cells to 7.3% of reads in BCL2L1-i treated cells, but enriched to 66.7% of reads in BCL2L1-i and MCL1-i co-treated cells. Dots show n = 2 biological replicates. (L) Alleles containing E325K edits were highly depleted in BCL2L1-i-treated cells, whereas the triple mutant E322K, E325K, G326N allele was most enriched in BCL2L1-i and MCL1-i co-treated cells.

**Figure S5.**
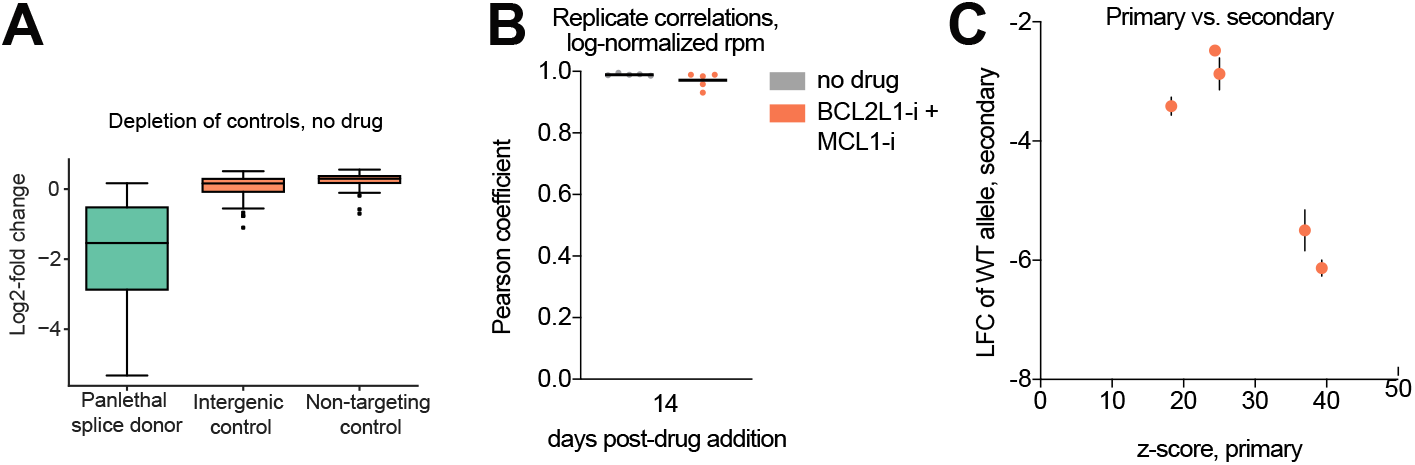
Quality control metrics for *BCL2L1* screen and validation experiments, related to Figure 5. (A) Distribution of log-fold change values of positive control sgRNAs (n = 32 essential splice site sgRNAs) and negative control sgRNAs (n = 75 intergenic sgRNAs, and n = 75 non-targeting sgRNAs). Boxes show the quartiles; whiskers show 1.5 times the interquartile range. (B) Distribution of Pearson coefficients for allele-level replicate correlations for sg24-sg28 at 14 days post-drug addition, either in the absence of drug (gray) or in the presence of BCL2L1-i and MCL1-i (orange). Only alleles with > 100 reads in at least one condition were considered. The mean is shown in black. (C) Comparison between sgRNA performance in the primary screen (z-score) and secondary validation (log fold change (LFC) of the wild-type allele at 14 days after addition of BCL2L1-i + MCl1-i). Error bars show the range of 2 biological replicates in the secondary validation.

**Figure S6.**
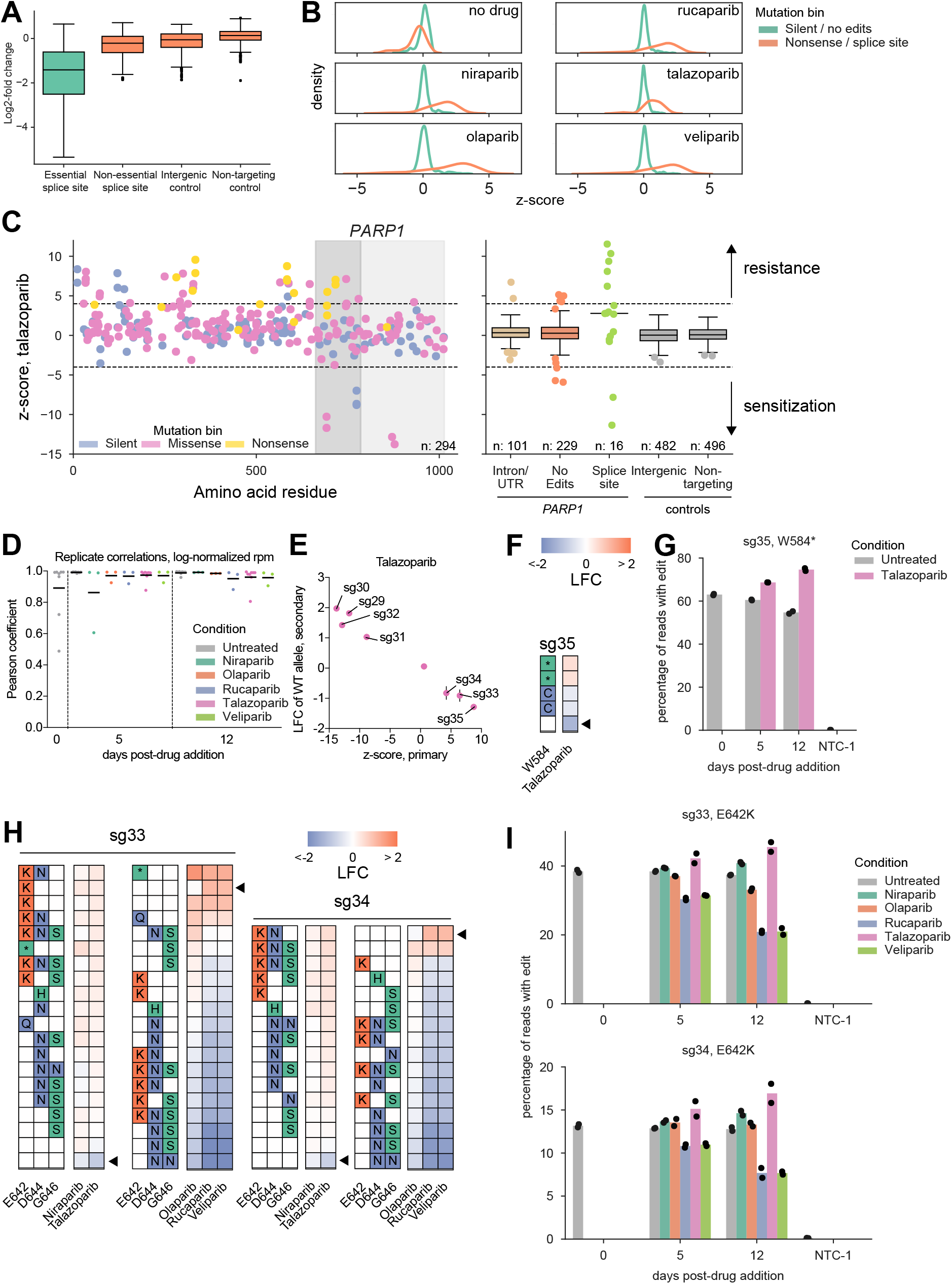
Mutagenesis of *PARP1* reveals missense mutations in the catalytic domain that sensitize to PARP inhibition, related to **Figure 6**. (A) Distribution of log-fold change values of positive control sgRNAs (n = 500 essential splice site sgRNAs) and negative control sgRNAs (n = 500 non-essential splice site sgRNAs, n = 482 intergenic sgRNAs, and n = 500 non-targeting sgRNAs). Boxes show the quartiles; whiskers show 1.5 times the interquartile range. (B) Distribution of sgRNAs predicted to introduce silent mutations or no edits (green) and sgRNAs predicted to introduce nonsense or splice site mutations (orange) in each drug condition. (C) Performance of sgRNAs targeting *PARP1* in the presence of talazoparib. Guides are colored and grouped according to the predicted mutation bin. Dashed lines show z-score thresholds of −4 (for depletion) and 4 (for enrichment); the HD domain and ART domain are shaded in dark gray and light gray, respectively. Boxes show the quartiles; whiskers show 1.5 times the interquartile range. Categories with n < 20 are shown as individual dots. (D) Distribution of Pearson coefficients for allele-level replicate correlations for sg29-sg36 at each timepoint and each drug condition. Only alleles with > 100 reads in at least one condition were considered. The mean is shown in black. (E) Comparison between sgRNA performance in the primary screen (z-score, talazoparib) and secondary validation (log-fold change (LFC) of the wild-type allele at 12 days post-talazoparib addition). Error bars show the range of 2 biological replicates in the secondary validation. Guides are labeled if the absolute LFC of the wild-type allele is > 0.5. (F) Alleles present in > 1% of reads in at least one replicate of sg35-treated cells. Amino acid changes for each allele are shown on the left; the average log-fold change of each allele (comparing talazoparib-treated to untreated cells at 12 days post-drug addition) is shown on the right. The wildtype (unedited) allele is marked with a black triangle. (G) Percentage of all sequencing reads containing the indicated edit for each timepoint and drug condition. Reads containing indels were not considered to be edited. NTC-1 indicates an untreated, non-targeting control (no sgRNA), sequenced at 12 days post-drug addition. Dots show n = 2 biological replicates. (H) Allele analysis, as in (F), for sg33- and sg34-treated sgRNAs in the presence of 5 PARPi. (I) Percentage of all sequencing reads containing the indicated edit for each timepoint and drug condition, as in (G), for sg33- and sg34-treated sgRNAs.

**Figure S7.**
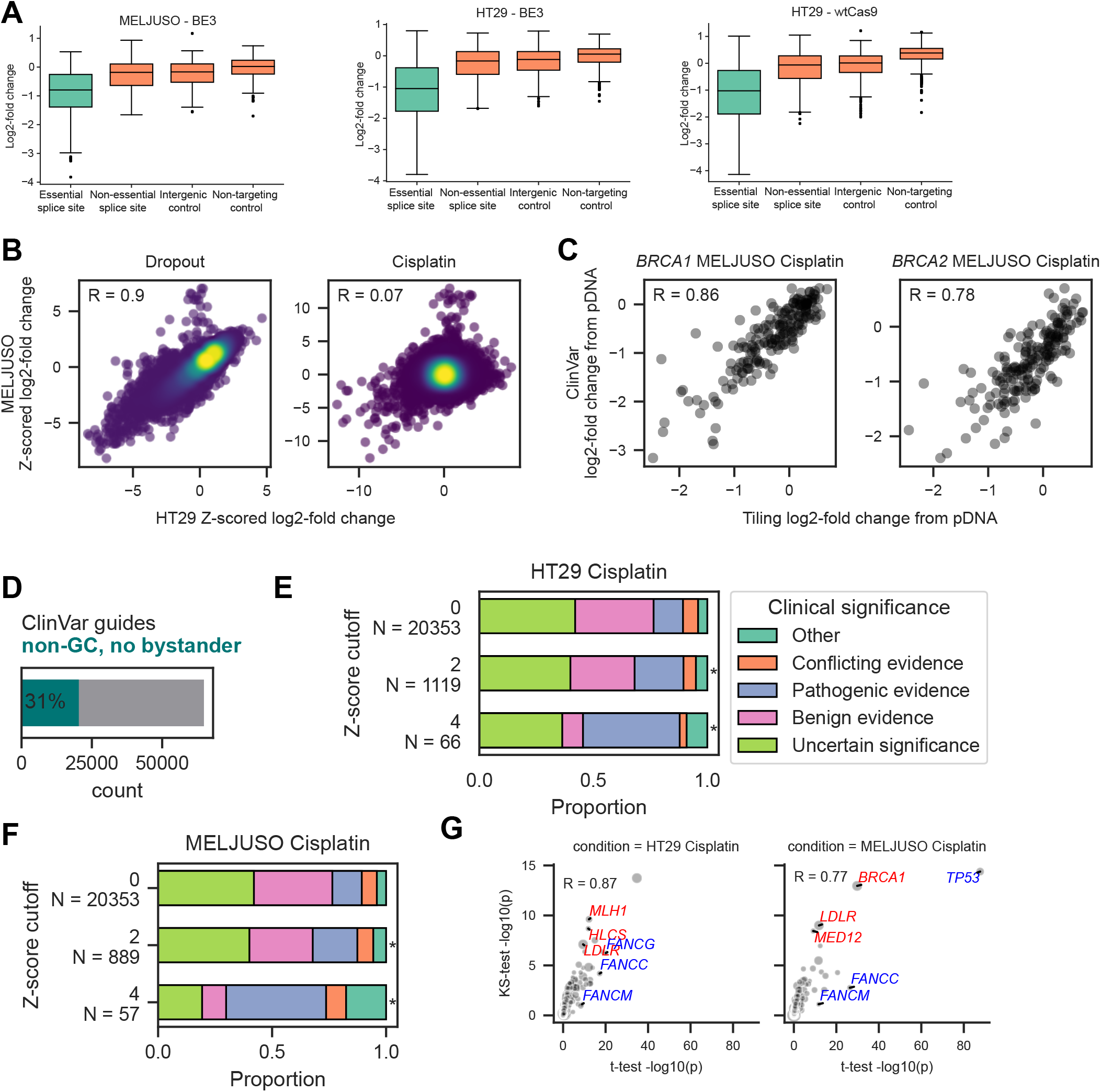
Profiling of 52,034 clinical variants using base editor, related to Figure 7. (A) Distribution of negative and positive control guides for ClinVar screens in MELJUSO and HT29 (1,461 negative and 496 positive controls) and wtCas9 in HT29 (1,456 negative and 496 positive controls) in the absence of small molecule treatment. Boxes show the quartiles; whiskers show 1.5 times the interquartile range. (B) Correlation of z-scored log-fold changes (relative to intergenic controls) between HT29 and MELJUSO dropout (left) and cisplatin (right) arms. Pearson correlation is indicated. Points are colored by density. (C) Correlation of log-fold changes of sgRNAs targeting BRCA1 (left) and BRCA2 (right) between ClinVar and prior tiling screens (Figure 2). Pearson correlation is indicated. (D) Fraction of guides predicted to make a “clean” edit (i.e. in a non-GC motif with no bystanders) in a clinical variant. (E) Distribution of clinical significance for guides predicted to make a clean edit at a range of Z-score cutoffs in HT29 treated with cisplatin. All cutoffs are significant when compared against the unfiltered distribution (*p < 1E-16, chi-squared test). (F) Same as (E) but for MELUSO treated with cisplatin. (G) Correlation of gene level significance between two statistical tests for HT29 and MELJUSO cisplatin arms. Top 3 genes by difference are labeled for the KS-test (red) and t-test (blue). Pearson correlation is indicated.

**Figure S8.**
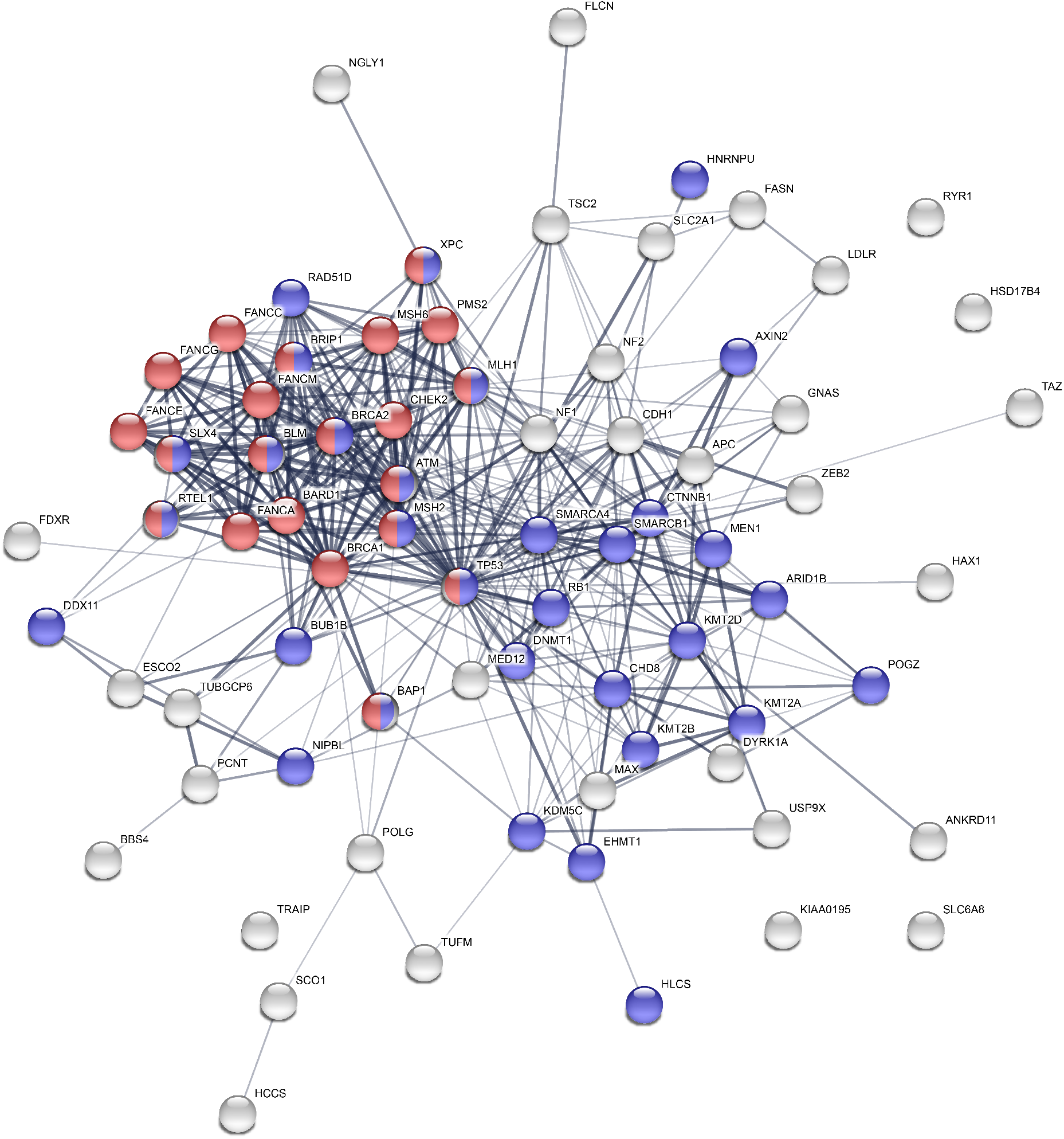
Protein-protein interactions between hits from HT29 and MELJUSO cisplatin screens, related to Figure 7. Edges represent confidence in STRING. Nodes represent gene hits. Nodes colored in red are in the Reactome DNA repair pathway. Nodes colored in blue are listed in the GO-term for chromatin organization. To define gene hits, we used an independent two sample t-test comparing the log-fold changes of VEP high impact sgRNAs to both intergenic and non-targeting controls in HT29 and MELJUSO treated with cisplatin. We then used the Benjamini-Hochberg procedure to define significant genes, using an FDR cutoff of 0.1. For each cell line, we identified hits which were significant when compared against both sets of controls. We then included all hits from either cell line in the network.

**Figure S9.**
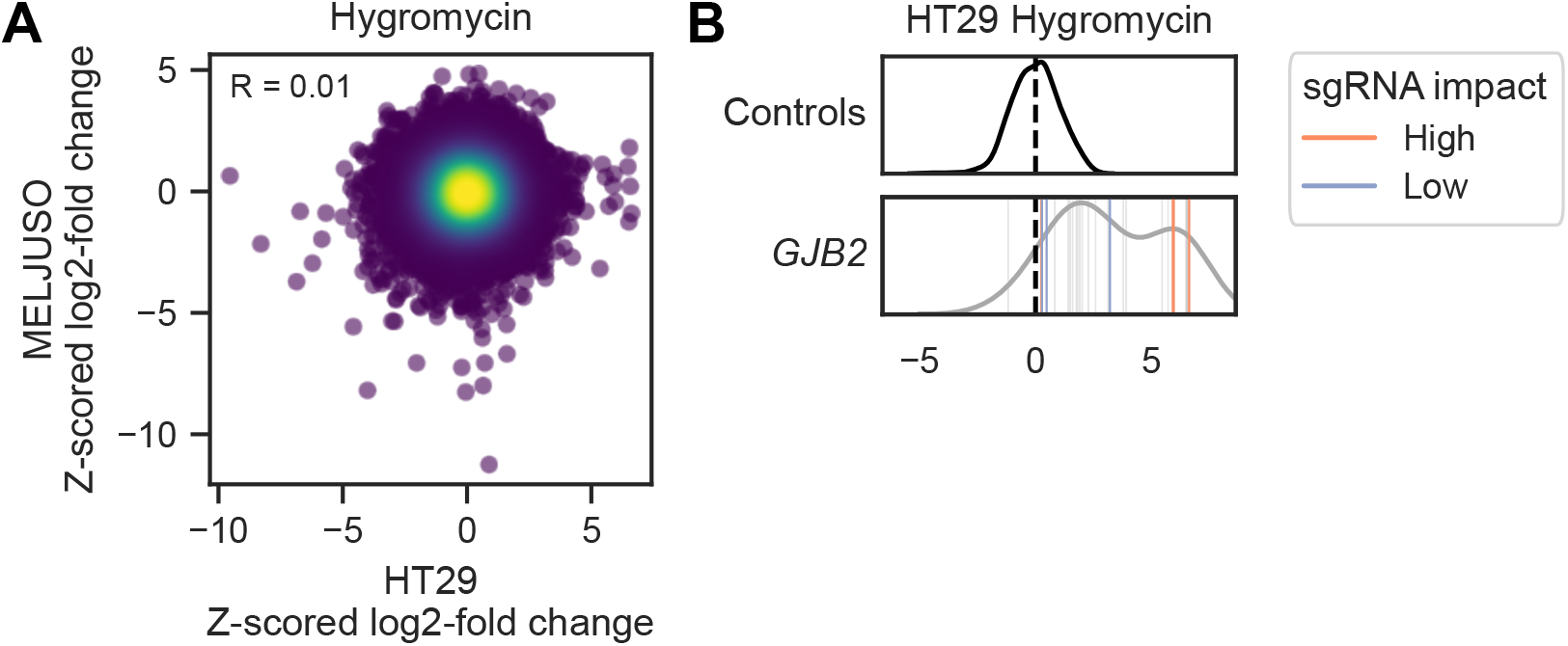
GJB2 loss-of-function in HT29 cells treated with hygromycin, related to Figure 7. (A) Correlation of z-scored log-fold changes (relative to intergenic controls) between HT29 and MELJUSO hygromycin arms. Pearson correlation is indicated. Points are colored by density. (B) Z-scored log-fold change for GJB2 in HT29 treated with hygromycin. The density plot for intergenic controls is plotted. The mean value for controls is indicated as a dashed line. Each bar represents a guide, and the density plot represents the distribution of guides targeting GJB2.

**Figure S10.**
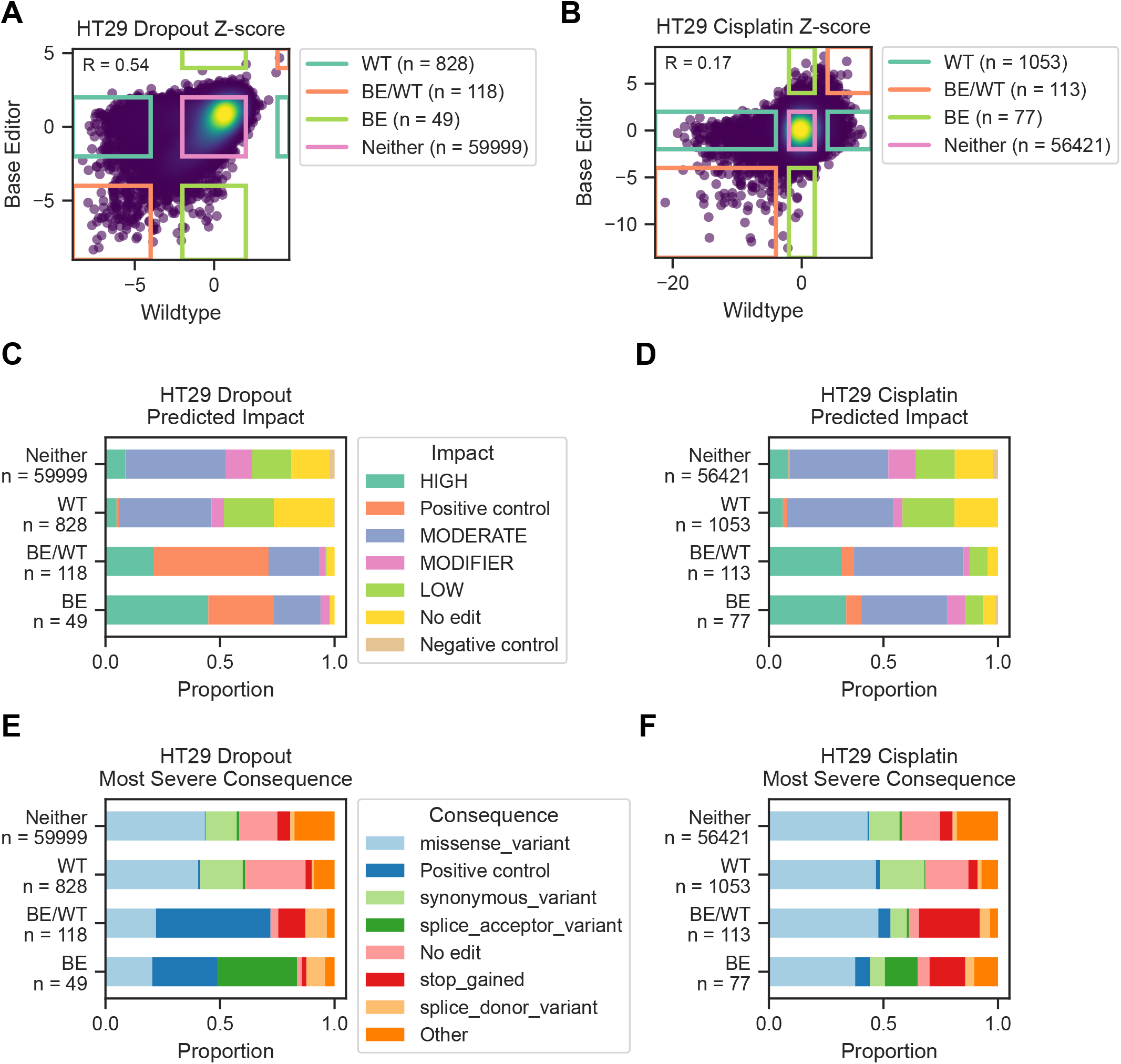
Counter-screen with wtCas9 in HT29 cells confirms loss-of-function alleles and uncovers potential gain-of-function mutants, related to Figure 7. (A) Correlation of z-scored log-fold changes (relative to intergenic controls) between wildtype (WT) and base editor (BE) Cas9 in the HT29 untreated arm. Bins for defining hits are outlined. Pearson correlation coefficient is indicated. Points are colored by density (B). Same as (A) but for HT29 treated with cisplatin. (C) Distribution of predicted impact for sgRNAs in the HT29 untreated arm. Bins correspond to those indicated in panel (A). Positive controls include sgRNAs targeting essential splice sites. Negative controls include sgRNAs targeting non-essential splice sites, intergenic and non-targeting controls. “No edit” sgRNAs only have C’s in a GC-motif in the edit window. (D) Same as (C) but for HT29 treated with cisplatin. (E) Same as (C) but for predicted consequence. All consequences with a maximum fraction of 0.05 across bins are grouped as “other”. (F) Same as (E) but for HT29 treated with cisplatin.

## REFERENCES

Beale, R.C.L., Petersen-Mahrt, S.K., Watt, I.N., Harris, R.S., Rada, C., and Neuberger, M.S. (2004). Comparison of the differential context-dependence of DNA deamination by APOBEC enzymes: correlation with mutation spectra in vivo. J. Mol. Biol. 337, 585–596.

Billon, P., Bryant, E.E., Joseph, S.A., Nambiar, T.S., Hayward, S.B., Rothstein, R., and Ciccia, A. (2017). CRISPR-Mediated Base Editing Enables Efficient Disruption of Eukaryotic Genes through Induction of STOP Codons. Mol. Cell 67, 1068–1079.e4.

Chang, W.-C., Wang, H.-C., Cheng, W.-C., Yang, J.-C., Chung, W.-M., Ho, Y.-P., Chen, L., Hung, Y.-C., and Ma, W.-L. (2020). LDLR-mediated lipidome-transcriptome reprogramming in cisplatin insensitivity. Endocr. Relat. Cancer 27, 81–95.

Chen, H., Liu, S., Padula, S., Lesman, D., Griswold, K., Lin, A., Zhao, T., Marshall, J.L., and Chen, F. (2020). Efficient, continuous mutagenesis in human cells using a pseudo-random DNA editor. Nat. Biotechnol. 38, 165–168.

Chen, H.-Y., Lang, Y.-D., Lin, H.-N., Liu, Y.-R., Liao, C.-C., Nana, A.W., Yen, Y., and Chen, R.-H. (2019). miR-103/107 prolong Wnt/β-catenin signaling and colorectal cancer stemness by targeting Axin2. Sci. Rep. 9, 9687.

Chou, C.-H., Lee, R.-S., and Yang-Yen, H.-F. (2006). An internal EELD domain facilitates mitochondrial targeting of Mcl-1 via a Tom70-dependent pathway. Mol. Biol. Cell 17, 3952–3963.

Clement, K., Rees, H., Canver, M.C., Gehrke, J.M., Farouni, R., Hsu, J.Y., Cole, M.A., Liu, D.R., Joung, J.K., Bauer, D.E., et al. (2019). CRISPResso2 provides accurate and rapid genome editing sequence analysis. Nat. Biotechnol. 37, 224–226.

Clohessy, J.G., and Zhuang, J. (2004). Characterisation of Mcl-1 cleavage during apoptosis of haematopoietic cells. British Journal of.

Cunningham, F., Achuthan, P., Akanni, W., Allen, J., Amode, M.R., Armean, I.M., Bennett, R., Bhai, J., Billis, K., Boddu, S., et al. (2019). Ensembl 2019. Nucleic Acids Res. 47, D745–D751.

Datlinger, P., Rendeiro, A.F., Schmidl, C., Krausgruber, T., Traxler, P., Klughammer, J., Schuster, L.C., Kuchler, A., Alpar, D., and Bock, C. (2017). Pooled CRISPR screening with single-cell transcriptome readout. Nat. Methods 14, 297–301.

Dawicki-McKenna, J.M., Langelier, M.-F., DeNizio, J.E., Riccio, A.A., Cao, C.D., Karch, K.R., McCauley, M., Steffen, J.D., Black, B.E., and Pascal, J.M. (2015). PARP-1 Activation Requires Local Unfolding of an Autoinhibitory Domain. Mol. Cell 60, 755–768.

Despres, P.C., Dubé, A.K., Seki, M., Yachie, N., and Landry, C.R. (2020). Perturbing proteomes at single residue resolution using base editing. Nat. Commun. 11, 1871.

Dixit, A., Parnas, O., Li, B., Chen, J., Fulco, C.P., Jerby-Arnon, L., Marjanovic, N.D., Dionne, D., Burks, T., Raychowdhury, R., et al. (2016). Perturb-Seq: Dissecting Molecular Circuits with Scalable Single-Cell RNA Profiling of Pooled Genetic Screens. Cell 167, 1853–1866.e17.

Doench, J.G., Fusi, N., Sullender, M., Hegde, M., Vaimberg, E.W., Donovan, K.F., Smith, I., Tothova, Z., Wilen, C., Orchard, R., et al. (2016). Optimized sgRNA design to maximize activity and minimize off-target effects of CRISPR-Cas9. Nat. Biotechnol. 34, 184–191.

Donovan, K.F., Hegde, M., Sullender, M., Vaimberg, E.W., Johannessen, C.M., Root, D.E., and Doench, J.G. (2017). Creation of Novel Protein Variants with CRISPR/Cas9-Mediated Mutagenesis: Turning a Screening By-Product into a Discovery Tool. PLoS One 12, e0170445.

Farmer, H., McCabe, N., Lord, C.J., Tutt, A.N.J., Johnson, D.A., Richardson, T.B., Santarosa, M., Dillon, K.J., Hickson, I., Knights, C., et al. (2005). Targeting the DNA repair defect in BRCA mutant cells as a therapeutic strategy. Nature 434, 917–921.

Feldman, D., Singh, A., Schmid-Burgk, J.L., Carlson, R.J., Mezger, A., Garrity, A.J., Zhang, F., and Blainey, P.C. (2019). Optical Pooled Screens in Human Cells. Cell 179, 787–799.e17.

Findlay, G.M., Boyle, E.A., Hause, R.J., Klein, J.C., and Shendure, J. (2014). Saturation editing of genomic regions by multiplex homology-directed repair. Nature 513, 120–123.

Findlay, G.M., Daza, R.M., Martin, B., Zhang, M.D., Leith, A.P., Gasperini, M., Janizek, J.D., Huang, X., Starita, L.M., and Shendure, J. (2018). Accurate classification of BRCA1 variants with saturation genome editing. Nature 562, 217–222.

Gaudelli, N.M., Komor, A.C., Rees, H.A., Packer, M.S., Badran, A.H., Bryson, D.I., and Liu, D.R. (2017). Programmable base editing of A•T to G•C in genomic DNA without DNA cleavage. Nature 551, 464–471.

Gaudelli, N.M., Lam, D.K., Rees, H.A., Solá-Esteves, N.M., Barrera, L.A., Born, D.A., Edwards, A., Gehrke, J.M., Lee, S.-J., Liquori, A.J., et al. (2020). Directed evolution of adenine base editors with increased activity and therapeutic application. Nat. Biotechnol.

Gelman, H., Dines, J.N., Berg, J., Berger, A.H., Brnich, S., Hisama, F.M., James, R.G., Rubin, A.F., Shendure, J., Shirts, B., et al. (2019). Recommendations for the collection and use of multiplexed functional data for clinical variant interpretation. Genome Med. 11, 85.

Hamilton, E.M., Polder, E., Vanderver, A., Naidu, S., Schiffmann, R., Fisher, K., Raguž, A.B., Blumkin, L., H-ABC Research Group, van Berkel, C.G.M., et al. (2014). Hypomyelination with atrophy of the basal ganglia and cerebellum: further delineation of the phenotype and genotype-phenotype correlation. Brain 137, 1921–1930.

Han, K., Jeng, E.E., Hess, G.T., Morgens, D.W., Li, A., and Bassik, M.C. (2017). Synergistic drug combinations for cancer identified in a CRISPR screen for pairwise genetic interactions. Nat. Biotechnol. 35, 463–474.

Hart, T., Brown, K.R., Sircoulomb, F., Rottapel, R., and Moffat, J. (2014). Measuring error rates in genomic perturbation screens: gold standards for human functional genomics. Mol. Syst. Biol. 10, 733.

Hart, T., Chandrashekhar, M., Aregger, M., Steinhart, Z., Brown, K.R., MacLeod, G., Mis, M., Zimmermann, M., Fradet-Turcotte, A., Sun, S., et al. (2015). High-Resolution CRISPR Screens Reveal Fitness Genes and Genotype-Specific Cancer Liabilities. Cell 163, 1515–1526.

Herrant, M., Jacquel, A., Marchetti, S., Belhacène, N., Colosetti, P., Luciano, F., and Auberger, P. (2004). Cleavage of Mcl-1 by caspases impaired its ability to counteract Bim-induced apoptosis. Oncogene 23, 7863–7873.

Hess, G.T., Frésard, L., Han, K., Lee, C.H., Li, A., Cimprich, K.A., Montgomery, S.B., and Bassik, M.C. (2016). Directed evolution using dCas9-targeted somatic hypermutation in mammalian cells. Nat. Methods 13, 1036–1042.

Hill, A.J., McFaline-Figueroa, J.L., Starita, L.M., Gasperini, M.J., Matreyek, K.A., Packer, J., Jackson, D., Shendure, J., and Trapnell, C. (2018). On the design of CRISPR-based single-cell molecular screens. Nat. Methods 15, 271–274.

Hillenmeyer, M.E., Fung, E., Wildenhain, J., Pierce, S.E., Hoon, S., Lee, W., Proctor, M., St Onge, R.P., Tyers, M., Koller, D., et al. (2008). The chemical genomic portrait of yeast: uncovering a phenotype for all genes. Science 320, 362–365.

Horlbeck, M.A., Xu, A., Wang, M., Bennett, N.K., Park, C.Y., Bogdanoff, D., Adamson, B., Chow, E.D., Kampmann, M., Peterson, T.R., et al. (2018). Mapping the Genetic Landscape of Human Cells. Cell 174, 953–967.e22.

Jun, S., Lim, H., Chun, H., Lee, J.H., and Bang, D. (2020). Single-cell analysis of a mutant library generated using CRISPR-guided deaminase in human melanoma cells. Commun Biol 3, 154.

Kelsell, D.P., Dunlop, J., Stevens, H.P., Lench, N.J., Liang, J.N., Parry, G., Mueller, R.F., and Leigh, I.M. (1997). Connexin 26 mutations in hereditary non-syndromic sensorineural deafness. Nature 387, 80–83.

Kim, Y.B., Komor, A.C., Levy, J.M., Packer, M.S., Zhao, K.T., and Liu, D.R. (2017). Increasing the genome-targeting scope and precision of base editing with engineered Cas9-cytidine deaminase fusions. Nat. Biotechnol. 35, 371–376.

Kleinstiver, B.P., Sousa, A.A., Walton, R.T., Tak, Y.E., Hsu, J.Y., Clement, K., Welch, M.M., Horng, J.E., Malagon-Lopez, J., Scarfò, I., et al. (2019). Engineered CRISPR-Cas12a variants with increased activities and improved targeting ranges for gene, epigenetic and base editing. Nat. Biotechnol.

Kluesner, M.G., Lahr, W.S., Lonetree, C.-L., Smeester, B.A., Claudio-Vázquez, P.N., Pitzen, S.P., Vignes, M.J., Lee, S.C., Bingea, S.P., Andrew, A.A., et al. (2020). CRISPR-Cas9 cytidine and adenosine base editing of splice-sites mediates highly-efficient disruption of proteins in primary cells.

Koblan, L.W., Doman, J.L., Wilson, C., Levy, J.M., Tay, T., Newby, G.A., Maianti, J.P., Raguram, A., and Liu, D.R. (2018). Improving cytidine and adenine base editors by expression optimization and ancestral reconstruction. Nat. Biotechnol. 36, 843–846.

Komor, A.C., Kim, Y.B., Packer, M.S., Zuris, J.A., and Liu, D.R. (2016). Programmable editing of a target base in genomic DNA without double-stranded DNA cleavage. Nature 533, 420.

Kotschy, A., Szlavik, Z., Murray, J., Davidson, J., Maragno, A.L., Le Toumelin-Braizat, G., Chanrion, M., Kelly, G.L., Gong, J.-N., Moujalled, D.M., et al. (2016). The MCL1 inhibitor S63845 is tolerable and effective in diverse cancer models. Nature 538, 477–482.

Kuscu, C., Parlak, M., Tufan, T., Yang, J., Szlachta, K., Wei, X., Mammadov, R., and Adli, M. (2017). CRISPR-STOP: gene silencing through base-editing-induced nonsense mutations. Nat. Methods 14, 710–712.

Kweon, J., Jang, A.-H., Shin, H.R., See, J.-E., Lee, W., Lee, J.W., Chang, S., Kim, K., and Kim, Y. (2019). A CRISPR-based base-editing screen for the functional assessment of BRCA1 variants. Oncogene.

Landrum, M.J., Lee, J.M., Benson, M., Brown, G.R., Chao, C., Chitipiralla, S., Gu, B., Hart, J., Hoffman, D., Jang, W., et al. (2018). ClinVar: improving access to variant interpretations and supporting evidence. Nucleic Acids Res. 46, D1062–D1067.

Leverson, J.D., Phillips, D.C., Mitten, M.J., Boghaert, E.R., Diaz, D., Tahir, S.K., Belmont, L.D., Nimmer, P., Xiao, Y., Ma, X.M., et al. (2015). Exploiting selective BCL-2 family inhibitors to dissect cell survival dependencies and define improved strategies for cancer therapy. Sci. Transl. Med. 7, 279ra40.

Levy, J.M., Yeh, W.-H., Pendse, N., Davis, J.R., Hennessey, E., Butcher, R., Koblan, L.W., Comander, J., Liu, Q., and Liu, D.R. (2020). Cytosine and adenine base editing of the brain, liver, retina, heart and skeletal muscle of mice via adeno-associated viruses. Nat Biomed Eng 4, 97–110.

Ma, Y., Zhang, J., Yin, W., Zhang, Z., Song, Y., and Chang, X. (2016). Targeted AID-mediated mutagenesis (TAM) enables efficient genomic diversification in mammalian cells. Nat. Methods 13, 1029–1035.

McLaren, W., Gil, L., Hunt, S.E., Riat, H.S., Ritchie, G.R.S., Thormann, A., Flicek, P., and Cunningham, F. (2016). The Ensembl Variant Effect Predictor. Genome Biol. 17, 122.

Mehta, I.S., Kulashreshtha, M., Chakraborty, S., Kolthur-Seetharam, U., and Rao, B.J. (2013). Chromosome territories reposition during DNA damage-repair response. Genome Biol. 14, R135.

Melnikov, A., Rogov, P., Wang, L., Gnirke, A., and Mikkelsen, T.S. (2014). Comprehensive mutational scanning of a kinase in vivo reveals substrate-dependent fitness landscapes. Nucleic Acids Res. 42, e112.

Miller, S.M., Wang, T., Randolph, P.B., Arbab, M., Shen, M.W., Huang, T.P., Matuszek, Z., Newby, G.A., Rees, H.A., and Liu, D.R. (2020). Continuous evolution of SpCas9 variants compatible with non-G PAMs. Nat. Biotechnol.

Najm, F.J., Strand, C., Donovan, K.F., Hegde, M., Sanson, K.R., Vaimberg, E.W., Sullender, M.E., Hartenian, E., Kalani, Z., Fusi, N., et al. (2017). Orthologous CRISPR-Cas9 enzymes for combinatorial genetic screens. Nat. Biotechnol. 36, 179.

Neggers, J.E., Kwanten, B., Dierckx, T., Noguchi, H., Voet, A., Bral, L., Minner, K., Massant, B., Kint, N., Delforge, M., et al. (2018). Target identification of small molecules using large-scale CRISPR-Cas mutagenesis scanning of essential genes. Nat. Commun. 9, 502.

Nishimasu, H., Shi, X., Ishiguro, S., Gao, L., Hirano, S., Okazaki, S., Noda, T., Abudayyeh, O.O., Gootenberg, J.S., Mori, H., et al. (2018). Engineered CRISPR-Cas9 nuclease with expanded targeting space. Science 361, 1259–1262.

Patwardhan, R.P., Lee, C., Litvin, O., Young, D.L., Pe’er, D., and Shendure, J. (2009). High-resolution analysis of DNA regulatory elements by synthetic saturation mutagenesis. Nat. Biotechnol. 27, 1173–1175.

Pettitt, S.J., Krastev, D.B., Brandsma, I., Dréan, A., Song, F., Aleksandrov, R., Harrell, M.I., Menon, M., Brough, R., Campbell, J., et al. (2018). Genome-wide and high-density CRISPR-Cas9 screens identify point mutations in PARP1 causing PARP inhibitor resistance. Nat. Commun. 9, 1849.

Rees, H.A., and Liu, D.R. (2018). Base editing: precision chemistry on the genome and transcriptome of living cells. Nat. Rev. Genet. 19, 770–788.

Replogle, J.M., Norman, T.M., Xu, A., Hussmann, J.A., Chen, J., Cogan, J.Z., Meer, E.J., Terry, J.M., Riordan, D.P., Srinivas, N., et al. (2020). Combinatorial single-cell CRISPR screens by direct guide RNA capture and targeted sequencing. Nat. Biotechnol.

Richter, M.F., Zhao, K.T., Eton, E., Lapinaite, A., Newby, G.A., Thuronyi, B.W., Wilson, C., Koblan, L.W., Zeng, J., Bauer, D.E., et al. (2020). Phage-assisted evolution of an adenine base editor with improved Cas domain compatibility and activity. Nat. Biotechnol. 1–9.

Ruffner, H., Joazeiro, C.A., Hemmati, D., Hunter, T., and Verma, I.M. (2001). Cancer-predisposing mutations within the RING domain of BRCA1: loss of ubiquitin protein ligase activity and protection from radiation hypersensitivity. Proc. Natl. Acad. Sci. U. S. A. 98, 5134–5139.

Sanson, K.R., DeWeirdt, P.C., Sangree, A.K., Hanna, R.E., Hegde, M., Teng, T., Borys, S.M., Strand, C., Keith Joung, J., Kleinstiver, B., et al. (2019). Optimization of AsCas12a for combinatorial genetic screens in human cells.

Shalem, O., Sanjana, N.E., Hartenian, E., Shi, X., Scott, D.A., Mikkelson, T., Heckl, D., Ebert, B.L., Root, D.E., Doench, J.G., et al. (2014). Genome-scale CRISPR-Cas9 knockout screening in human cells. Science 343, 84–87.

Sharon, E., Chen, S.-A.A., Khosla, N.M., Smith, J.D., Pritchard, J.K., and Fraser, H.B. (2018). Functional Genetic Variants Revealed by Massively Parallel Precise Genome Editing. Cell 175, 544–557.e16.

Starita, L.M., Ahituv, N., Dunham, M.J., Kitzman, J.O., Roth, F.P., Seelig, G., Shendure, J., and Fowler, D.M. (2017). Variant Interpretation: Functional Assays to the Rescue. Am. J. Hum. Genet. 101, 315–325.

Szklarczyk, D., Gable, A.L., Lyon, D., Junge, A., Wyder, S., Huerta-Cepas, J., Simonovic, M., Doncheva, N.T., Morris, J.H., Bork, P., et al. (2019). STRING v11: protein-protein association networks with increased coverage, supporting functional discovery in genome-wide experimental datasets. Nucleic Acids Res. 47, D607–D613.

Tao, Z.-F., Hasvold, L., Wang, L., Wang, X., Petros, A.M., Park, C.H., Boghaert, E.R., Catron, N.D., Chen, J., Colman, P.M., et al. (2014). Discovery of a Potent and Selective BCL-XL Inhibitor with in Vivo Activity. ACS Med. Chem. Lett. 5, 1088–1093.

Vinyard, M.E., Su, C., Siegenfeld, A.P., Waterbury, A.L., Freedy, A.M., Gosavi, P.M., Park, Y., Kwan, E.E., Senzer, B.D., Doench, J.G., et al. (2019). CRISPR-suppressor scanning reveals a nonenzymatic role of LSD1 in AML. Nat. Chem. Biol. 15, 529–539.

Walton, R.T., Christie, K.A., Whittaker, M.N., and Kleinstiver, B.P. (2020). Unconstrained genome targeting with near-PAMIess engineered CRISPR-Cas9 variants. Science 368, 290–296.

Weile, J., Sun, S., Cote, A.G., Knapp, J., Verby, M., Mellor, J.C., Wu, Y., Pons, C., Wong, C., van Lieshout, N., et al. (2017). A framework for exhaustively mapping functional missense variants. Mol. Syst. Biol. 13, 957.

Yao, J., Huang, T., Fang, X., Chi, Y., Zhu, Y., Wan, Y., Matsue, H., and Kitamura, M. (2010). Disruption of gap junctions attenuates aminoglycoside-elicited renal tubular cell injury. Br. J. Pharmacol. 160, 2055–2068.

Yeh, W.-H., Chiang, H., Rees, H.A., Edge, A.S.B., and Liu, D.R. (2018). In vivo base editing of post-mitotic sensory cells. Nat. Commun. 9, 2184.

Zandarashvili, L., Langelier, M.-F., Velagapudi, U.K., Hancock, M.A., Steffen, J.D., Billur, R., Hannan, Z.M., Wicks, A.J., Krastev, D.B., Pettitt, S.J., et al. (2020). Structural basis for allosteric PARP-1 retention on DNA breaks. Science 368.

Zhang, S., Samocha, K.E., Rivas, M.A., Karczewski, K.J., Daly, E., Schmandt, B., Neale, B.M., MacArthur, D.G., and Daly, M.J. (2018). Base-specific mutational intolerance near splice sites clarifies the role of nonessential splice nucleotides. Genome Res. 28, 968–974.

Zimmermann, M., Murina, O., Reijns, M.A.M., Agathanggelou, A., Challis, R., Tarnauskaitė, Ž., Muir, M., Fluteau, A., Aregger, M., McEwan, A., et al. (2018). CRISPR screens identify genomic ribonucleotides as a source of PARP-trapping lesions. Nature 559, 285–289.

